# Transposable elements mediate genetic effects altering the expression of nearby genes in colorectal cancer

**DOI:** 10.1101/2021.12.03.471093

**Authors:** Nikolaos M. R. Lykoskoufis, Evarist Planet, Halit Ongen, Didier Trono, Emmanouil T. Dermitzakis

**Affiliations:** Department of Genetic Medicine and Development, University of Geneva Medical School, 1211 Geneva, Switzerland; Institute for Genetics and Genomics in Geneva (iGE3), University of Geneva, 1211 Geneva, Switzerland; Swiss Institute of Bioinformatics, 1211 Geneva, Switzerland; School of Life Sciences, Ecole Polytechnique Fédérale de Lausanne (EPFL), 1015, Lausanne, Switzerland

## Abstract

Transposable elements (TEs) are interspersed repeats that contribute to more than half of the human genome, and TE-embedded regulatory sequences are increasingly recognized as major components of the human regulome. Perturbations of this system can contribute to tumorigenesis, but the impact of TEs on gene expression in cancer cells remains to be fully assessed. Here, we analyzed 275 normal colon and 276 colorectal cancer (CRC) samples from the SYSCOL colorectal cancer cohort and discovered 10,111 and 5,152 TE expression quantitative trait loci (eQTLs) in normal and tumor tissues, respectively. Amongst the latter, 376 were exclusive to CRC, likely driven by changes in methylation patterns. We identified that transcription factors are more enriched in tumor-specific TE-eQTLs than shared TE-eQTLs, indicating that TEs are more specifically regulated in tumor than normal. Using Bayesian Networks to assess the causal relationship between eQTL variants, TEs and genes, we identified that 1,758 TEs are mediators of genetic effect, altering the expression of 1,626 nearby genes significantly more in tumor compared to normal, of which 51 are cancer driver genes. We show that tumor-specific TE-eQTLs trigger the driver capability of TEs subsequently impacting expression of nearby genes. Collectively, our results highlight a global profile of a new class of cancer drivers, thereby enhancing our understanding of tumorigenesis and providing potential new candidate mechanisms for therapeutic target development.

## INTRODUCTION

Understanding the mechanisms underlying tumorigenesis has been one of the main research questions in cancer biology. While somatic mutations, chromosomal rearrangements and gene amplification are the three main hallmarks driving cancer progression, they are unable to provide a complete explanation of tumorigenesis. Recent discoveries have demonstrated that transposable elements (TEs) have contributed to the evolution of gene regulation and can alter the landscape of gene expression in development and disease [1-5]. Transposable elements (TEs) are interspersed repeats that contribute more than half of the human genome. TEs, more specifically TE-embedded regulatory sequences (TEeRS) are broadly active during the phases of genome reprogramming that occur in the germline and the early embryo, and then controlled by epigenetic mechanisms that still allow their finely orchestrated participation in biological events as diverse as brain development, immune responses, and metabolic control. The aberrant re-activation of TEeRSs is observed under certain conditions and disease states, notably cancer [6-8]. Transcription is defined by the coordinated activity of regulatory elements which are modulated by genetic variation. Thus, we speculate that transposable element expression is influenced by regulatory non-coding variants, also called expression Quantitative Trait Loci (eQTLs), known to contribute to the onset and progression of complex diseases like cancer [9, 10]. To build on this concept, we set out to analyze the interplay between regulatory variants (eQTLs), transposable elements and gene expression to characterize the genetic perturbation of TE and gene expression in cancer. To this end, we integrated genome-wide genotyping data (genotype array) and transcriptomic profiles (bulk RNA-sequencing) from the Systems Biology of Colorectal Cancer (SYSCOL) cohort comprising of 275 and 276 normal and tumor samples, respectively.

## RESULTS

### Quantifying transposable elements (TEs) and gene expression

To measure the expression of TEs in CRC, we examined transcriptomes obtained by RNA-seq from 275 normal and 276 CRC samples from the SYSCOL cohort [11]. We quantified TE and gene expression using an in-house curated TE annotation list originating from the *RepBase* database [12] that contains approximately 4.6 million individual TE loci. These annotations were merged with gene annotation from *ensembl* (v19). Filtering for uniquely mapped reads (**Methods**) to obtain robust estimates of TE expression resulted in 51,320 TEs and 17,360 genes (protein coding and lincRNAs). We observed that the majority of expressed TEs present in our dataset are SINEs (Alu and MIR), LINEs (L1 and L2) as well as different subfamilies of Long Terminal Repeats (LTRs) and DNA transposons. However, when we looked at the proportion of expressed TEs per subfamily, SVA and ERVK were most prominent (**Figure 1A, Supplementary figure 1**). Additionally, we used available data from Encode [13] and miRbase [14] to generate a list of regulatory regions and discovered that 13,656 expressed TEs overlapped with at least one previously identified regulatory element. We also discovered that expressed TEs are significantly enriched for most regulatory regions, except for enhancers, compared to non-expressed TEs (**Supplementary table 1**; **Figure 1B, Supplementary figure 2**) highlighting their potential role in gene expression regulation.

**Figure 1:**
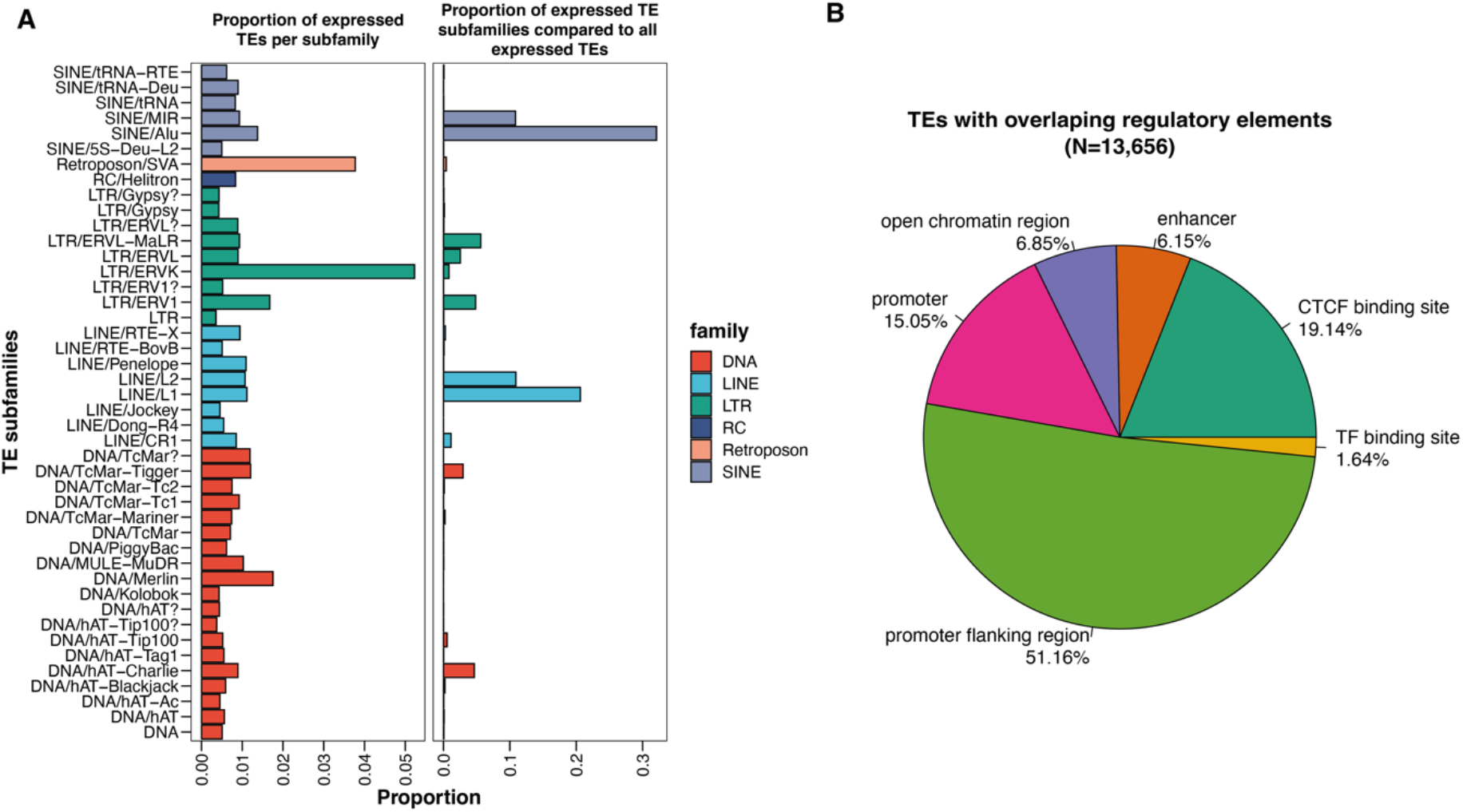
Description of quantified TEs. (**A**) Barplot showing the proportion of uniquely mapped and quantified TE subfamilies in our dataset. (**B**) Pie chart showing the proportion of TEs with different types of regulatory elements within their sequence. We uniquely mapped and quantified 51,320 TEs. The majority of them are SINEs from the Alu and MIR family, L1 and L2 TEs from the LINE family and different subfamilies of LTRs as well as some DNA transposons. When we looked at the proportion of expressed TE per subfamily, we observed that SVA and ERVK are most prominent. Additionally, 13,656 out of the 51,320 TEs contain regulatory elements within sequence.

### Transposable elements are under strong genetic control

Using TE expression quantifications and genotype data we first sought to assess the impact of inter-individual genetic variation on TE expression. We conducted *cis*-eQTL analysis followed by a forward backward stepwise conditional analysis (**Methods**) and discovered a total of 10’111 and 5’152 TE-eQTLs as well as 6’856 and 1’539 gene eQTLs in normal and tumoral tissue, respectively (**Supplementary Figure 3**,**4; Supplementary Tables 2**,**3**). Similarly to gene-eQTLs, TE-eQTLs displayed stronger effects and density closer to the transcription start site (TSS) in both normal and tumor samples (spearman rho=-0.34, P<2.2e-16 in normal, spearman rho=-0.27, P<2.2e-16 in tumor) (**Figure 2A-B**), yet were more proximal to the TSS compared to gene-eQTLs (Wilcoxon P=7.6e-11 in normal; Wilcoxon P=3.3e-05 in tumor; **Supplementary Figure 5**). We observed that TEs displayed fewer independent eQTLs per TE than genes (**Figure 2C-D**) while the minor allele frequencies of TE- and gene-eQTL variants were similar (**Supplementary Figure 6**). Proximal distance of TE-eQTLs to TSS and the smaller number of independent signals per TE could be due to smaller evolutionary time of TE regulatory landscapes in the human genome compared to genes, making proximal effects much more likely.

**Figure 2:**
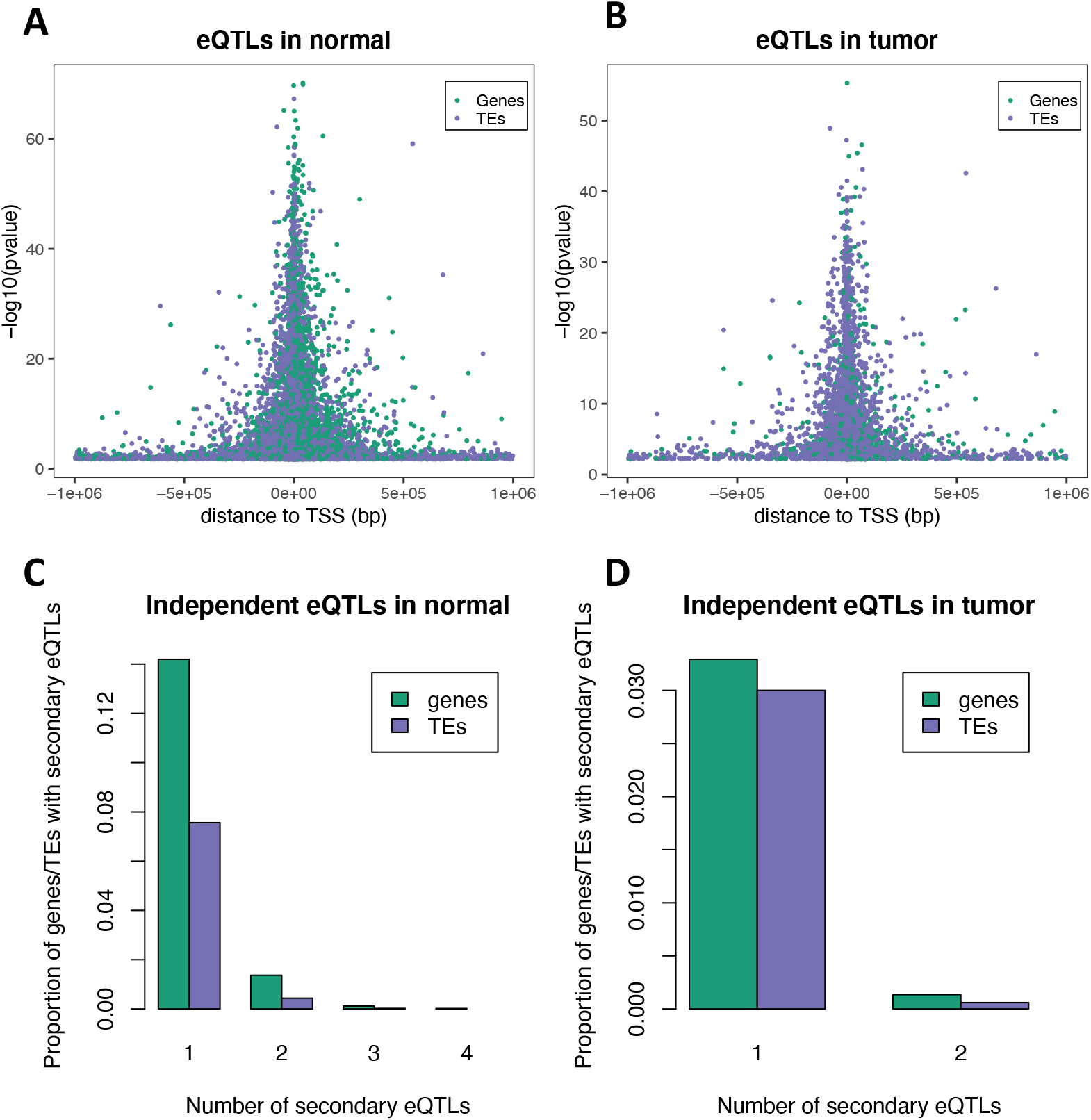
cis- eQTL discovery. eQTL variant distance to TSS in (A) normal and (B) in tumor. We observe stronger eQTL effect close to the transcription start site of TE and genes in both normal and tumor. Number of secondary eQTLs for TEs and genes in (C) normal and (D) tumor. Gene eQTLs have more functionally independent eQTLs per gene than TEs do.

Given previously established roles of tumor-specific gene-eQTLs in tumorigenesis [11], we aimed next at investigating whether tumor-specific TE-eQTLs could similarly contribute as cancer driving factors. To this end, we used linear mix models with an interaction term between variant and tissue (normal/tumor). We discovered that 376 (7.3%) of the tumor TE-eQTLs are tumor-specific and 1,685 (17%) of the normal TE-eQTLs are normal-specific, with 524 TE-eQTLs shared between both settings (**Figure 3A; Supplementary Table 5**). For genes, we found 101 (6.5%) tumor gene-eQTLs to be tumor-specific and 897 (13%) normal gene-eQTLs to be normal-specific, of which 169 were shared (**Supplementary Figure 7A; Supplementary Table 4**). Shared TE- and gene-eQTLs were closer to the TSS of TEs/genes compared to tissue-specific eQTLs (Wilcoxon P<2.2e-16) (**Figure 3B, Supplementary figure 7B)**. Additionally, we observed that shared eQTLs conserved their effect in both normal and tumor (**Figure 3C, Supplementary figure 7C**). These results indicate that TE expression is under strong genetic control and that non-coding germline variants act as drivers of TE expression in cancer as similarly observed for gene expression [11].

**Figure 3:**
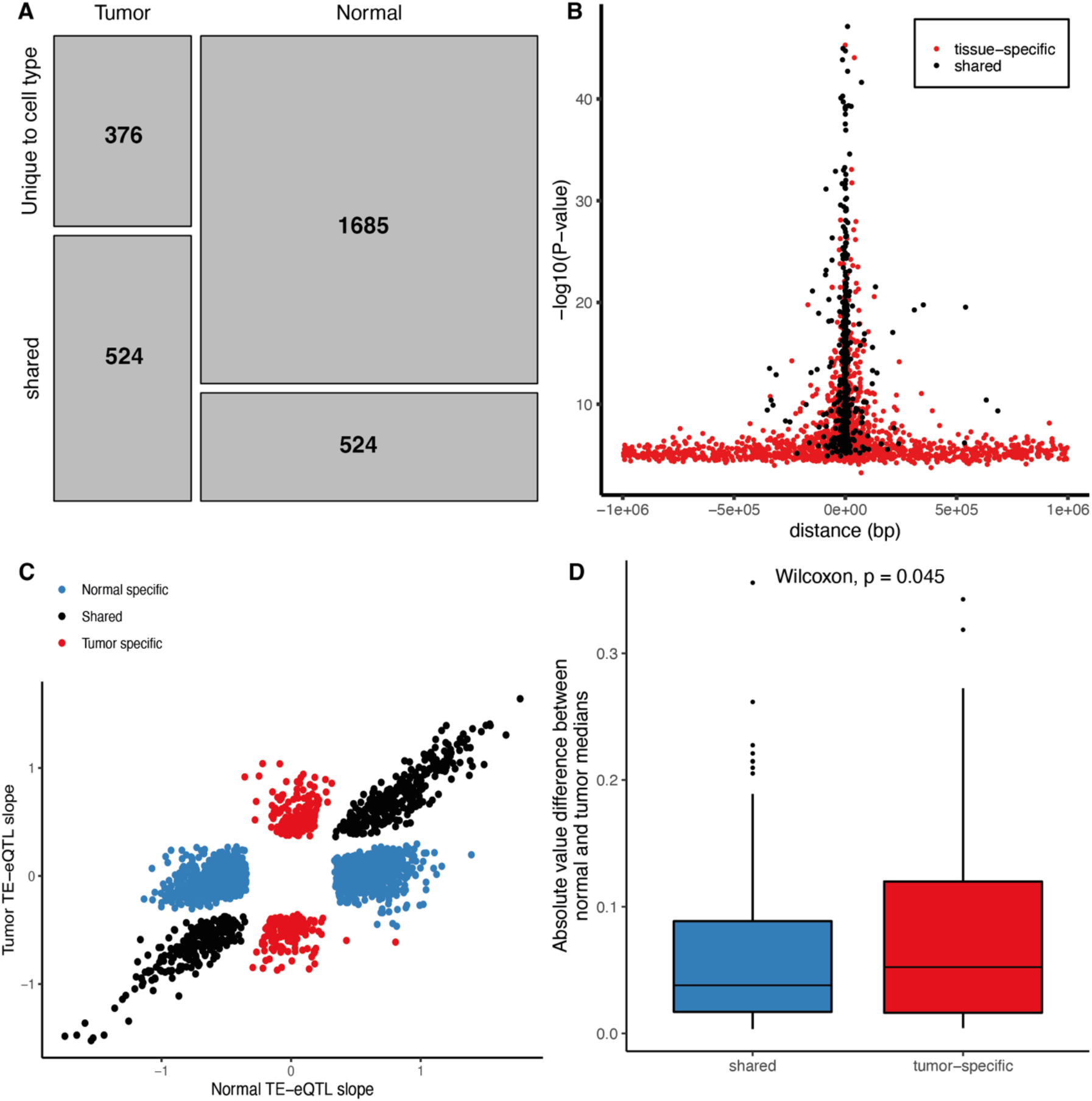
Tissue specificity of TE-eQTLs. (**A**) Mosaic plot of tissue specificity of TE-eQTLs. (**B**) Tissue specificity and distance of TE-eQTL to transcription start site (TSS). The shared TE-eQTLs (black) are closer to the TSS than are the tissue specific TE-eQTLs (red) (Wilcoxon P<2.2e-16). (**C**) TE-eQTL slopes for the normal specific TE-eQTLs in blue, the tumor specific in red and shared in black. (**D**) Boxplot of the absolute value difference of median methylation betas between normal and tumor samples for shared and tumor=specific TE-eQTLs.

### Transcription factors regulate TE expression more specifically in tumor

To corroborate the biological relevance of the discovered TE-eQTL variants we performed functional enrichment analysis of TE and gene eQTLs in normal and tumor using available ChIP-seq data from the Ensembl Regulatory Build [15] (**methods**). We found significant enrichment for many TF binding sites overlapping the eQTL loci highlighting the functional relevance of the variants discovered (**Figure 4A-B; Supplementary Figure 8**,**9; Supplementary Tables 6**,**7**). We discovered 5 TFs and 15 TFs that displayed stronger enrichment for TE eQTLs compared to gene eQTLs in normal and tumor, respectively. The TF most enriched over TE-eQTLs in normal tissues was ZNF274, a Krüppel-associated box (KRAB) domain-containing zinc-finger protein (KZFP), whereas the most enriched over tumor TE-eQTLs was TRIM28, the master corepressor that is recruited by the KRAB domain of many TE-binding KZFPs and serves as a scaffold for a heterochromatin-inducing complex capable of repressing TEs via histone H3 Lys9 trimethylation (H3K9me3), histone deacetylation and DNA methylation [16, 17]. Additionally, BDP1 and BRF1, two subunits of the RNA polymerase III transcription initiation factor, were more enriched over TE-eQTLs compared to gene eQTLs highlighting potential transcription of *Alu* or MIR TEs of the SINE family [18]. These results corroborate the biological relevance of TE eQTLs and point to possible transcription and repression of certain TEs.

**Figure 4:**
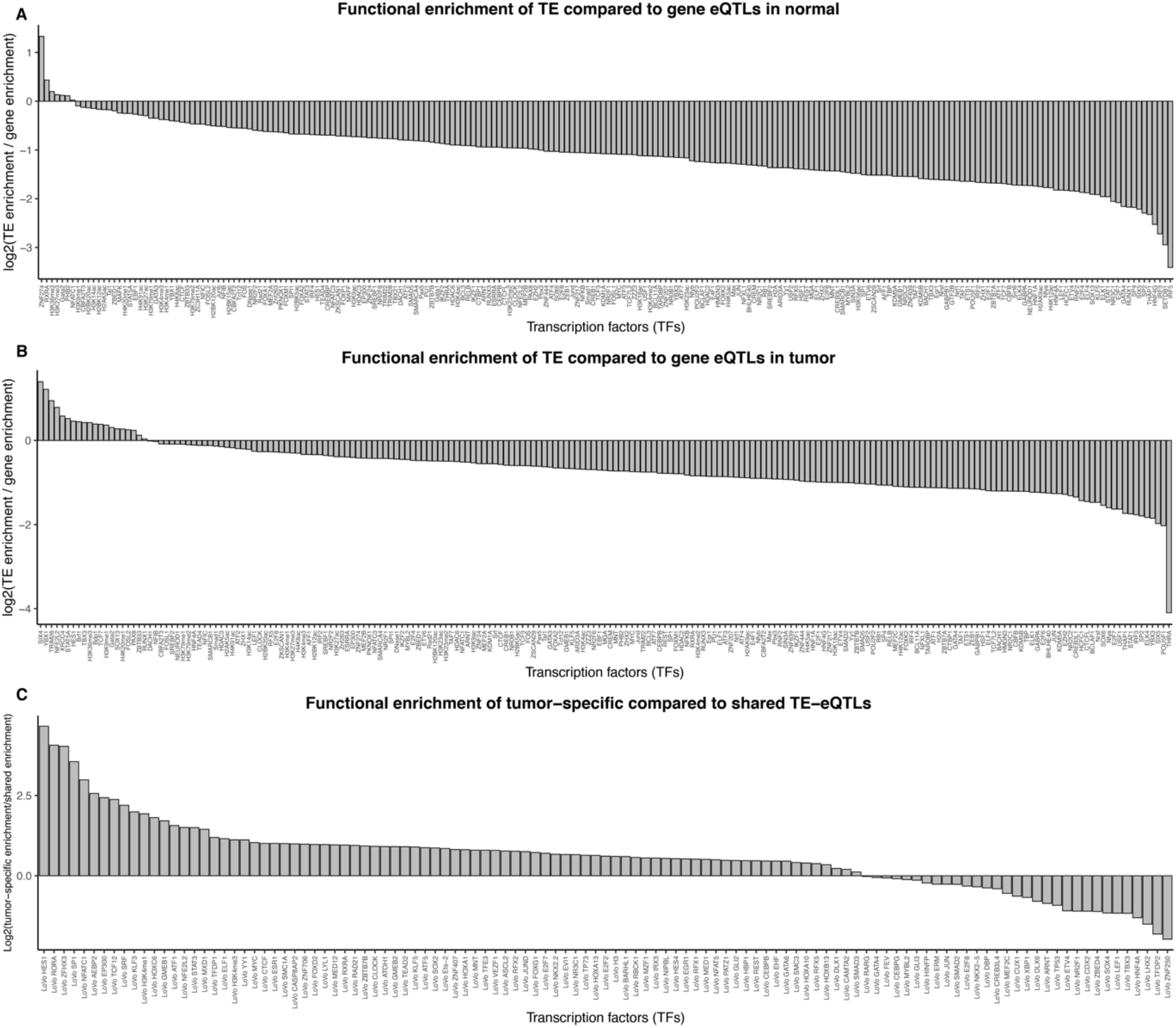
Functional enrichment of eQTLs. (A) The ratio between TE-eQTL enrichment and gene-eQTL enrichment in log2 scale discovered in normal. 5 TFs show stronger enrichment for TE-eQTLs in normal compared to gene-eQTLs. (B) The ratio between TE-eQTL enrichment and gene-eQTL enrichment in log2 scale discovered in tumor. We observed 15 TFs to have a stronger enrichment for TE-eQTLs than gene-eQTLs in normal. (C) log2 ratio between tumor-specific TE-eQTL enrichment and shared TE eQTL enrichment. We observe 80 TFs with a stronger enrichment for the tumor-specific TE-eQTLs than the shared eQTLs indicating that these TFs regulate TE expression specifically in tumor.

To assess the differential effects of tumor-specific versus shared eQTLs, we performed functional enrichment analyses using available ChIP-seq data from LoVo colorectal cancer cells [19] (**methods**). We observed that in the case of genes, all tested TFs had a stronger enrichment for shared compared to tumor-specific eQTLs, indicating that these TFs are regulating gene expression in both the normal and tumor state. (**Supplementary Figure 10, Supplementary Table 8**). In contrast, we found 80 TFs displaying stronger enrichment for tumor-specific versus shared TE-eQTLs, pointing to tumor-specific TE regulation (**Figure 4C; Supplementary Figure 11; Supplementary Table 9**). Of these, 39 were upregulated and 34 downregulated in tumors (7 were missing from our expression data and could not be tested for differential expression analysis), but we did not observe any significant correlation between the tumor-specific TE-eQTL enrichment to shared TE-eQTL enrichment ratio and fold change in the expression of the corresponding transcription factors (Pearson R = -0.18, p-value = 0.083; **Supplementary Figure 12**). Thus, differential expression of these TFs is not driving the tumor-specific TE-eQTL effects. However, 59 of the 86 tumor-specific TE-eQTLs overlapping the binding sites of the 80 aforementioned TFs are not significantly associated (FDR = 5%) with any nearby (±1 Mb from TSS) TE or gene in normal, indicating that these regulatory regions are inactive in the normal state (**Supplementary Figure 13**). Additionally, we compared methylation levels between normal and tumor samples for the tumor-specific and shared eQTLs and observed significantly increased (Wilcoxon rank sum test p-value = 0.017 for TEs and p-value = 0.00097 for genes) methylation over tumor-specific compared to shared eQTLs for both gene and TEs (**Figure 3D; Supplementary figure 7D**).

Altogether these results suggest that many TFs are regulating TE expression. The inactivity of some of the TE eQTLs in normal and the significant changes in methylation between tumor-specific and shared TE-eQTLs indicate that regulatory switches involving the recruitment of these TFs might underlie the effects of tumor-specific TE eQTLs.

### Transposable elements as mediators of genetic effects onto genes

Having established that TEs are under genetic control, we next sought to assess the causal relationship between eQTL variants, TEs and genes and discover the extent to which TEs act as drivers of gene expression in tumor. To this end, we focused on regulatory variants affecting both TEs and genes and detected these in an unbiased manner by first associating TEs with genes using a similar approach to QTL mapping. Next, we quantified the identified 21,263 TE-gene pairs found in normal samples and 144,289 TE-gene pairs found in tumor at 1% FDR and used this newly quantified TE-gene pairs to find all eQTL-TE-gene triplets by performing a standard eQTL analysis (**Methods; Supplementary figure 14-16**). At 5% FDR, we discovered 12,379 and 9,714 triplets in normal and tumor, respectively, for which we inferred the most likely causal relationship using Bayesian networks (**Methods**) [20-22]. We tested three models, (i) the causal model where the eQTL variant affects TE expression and then gene expression, (ii) the reactive model where the eQTL variant affects gene expression and then TE expression and (iii) the independent model where the eQTL variant affects independently TE and gene expression (**Supplementary figure 17**). We observed significantly more causal models in tumor (47%) compared to normal (22%) (Fisher p-value <2e^-16^) indicating that TEs are causal for gene expression predominantly in tumor and to a lesser extent in normal (**Figure 5A, B; Supplementary figure 18; Supplementary Table 10**,**11**). We also show that the proportion of causal models correlated with the genomic position of the TE with respect to the gene; intronic TEs tended to be reacting to gene expression whereas TEs outside the gene body tended to be causal. Interestingly, there were significantly more causal scenarios when the eQTL variant lied within the TE, rather than outside (Fisher p-value < 2e^-16^) pinpointing to direct regulatory effects of the TE onto gene expression (**Supplementary figure 19**). Altogether, these results show that TEs are significantly more causal for changes in gene expression in tumor than in normal tissue.

**Figure 5:**
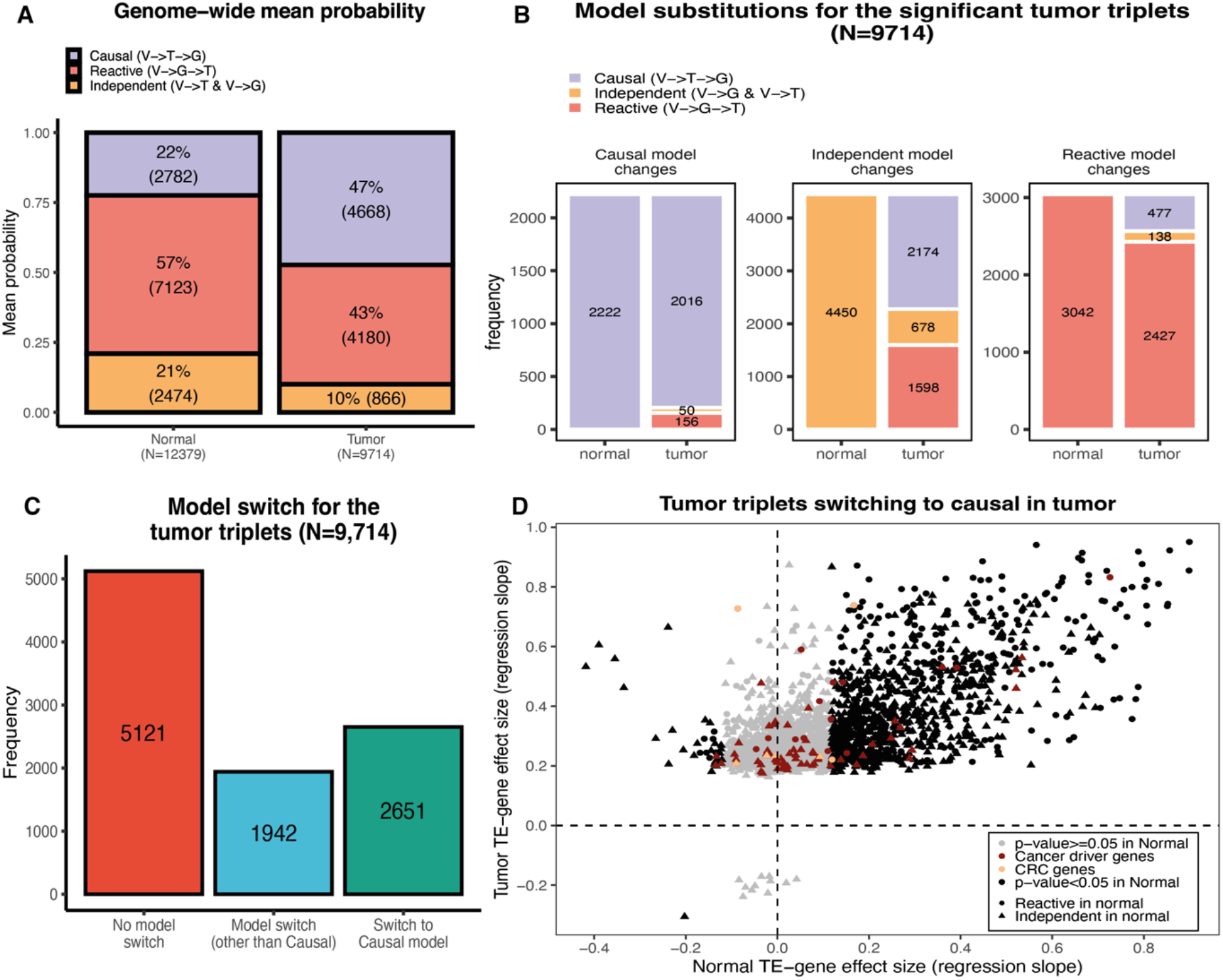
Causal relationship between eQTL variants, TEs and genes. (**A**) Barplot representing the mean probability for each of the three models in normal and tumor. We observe significantly more causal cases in tumor compared to normal (Wilcox P-value < 2e-16) (**B**) Barplot representing the model substitutions for the 9,714 tumor triplets from normal to tumor. Independent models tend to shift to a causal in tumor. This is true also for the reactive models in normal but to a much smaller extent. (**C**) Barplot representing the number of triplets that do not switch models, that switch to a causal model or that switch to reactive/independent from normal to tumor. The majority of triplets do not switch models between normal and tumor. However, 2,651 triplets are switching to a causal model making the corresponding TEs potential drivers of gene expression (**D**) Each point represents a TE-gene for each of the 2,561 tumor triplets. All points aresignificant in tumor but not in normal (grey points). We observe that in most cases, Tes are positively correlated with genes except for a few cases. Most cancer driver genes have no significant correlation with any TE in normal indicating that for most part, TEs impact them specifically in tumor.

### Transposable elements are drivers of gene expression during tumorigenesis

These results suggested that genetic variations in TE expression might drive tumorigenesis. To test this hypothesis, we considered the union of all triplets, i.e. the eQTL variant, TE and gene expression, discovered across tumor and normal tissue and using the same BN approach as previously mentioned, we inferred the causal relationship between the triplets in both states (**methods**). We similarly looked for shared triplets across the 12,379 normal and 9,714 tumor triplets (eQTL-TE-gene triplets are the same in both states or the eQTL for TE-gene pair is in high LD (R^2^ >=0.9)). In both shared and union triplets, we observed a significant increase in the causal model in tumor (Fisher Exact Test p-value < 2.2e^-16^ for shared and union triplets) mainly due to independent models and to a lesser extent reactive model shifting to causal. (**Supplementary figure 20; Supplementary Table 12**,**13**). Focusing on the 9,714 tumor triplets, we discovered 2,651 (28%) triplets that switched to a causal model in tumor compared to normal, highlighting regulatory changes whereby TEs impacted the expression of nearby genes (**Figure 5C**). These 2,651 triplets constituted of 1,758 unique TEs impacting 1,626 unique genes. Interestingly, we observed that TEs switching to causal were significantly up-regulated compared to TEs that did not switch models between normal and tumor or that switched but not to causal (Wilcoxon p-value 2.9e^-6^; **Supplementary Figure 21**). These results suggest that upregulation of TEs could give rise to their gene expression driver capability.

While expression of most TEs was positively correlated with the expression of the associated gene in tumor (n=2,639) (**Figure 5D**), only a few showed negative correlation (n=12). Of the significant tumor TE-gene pairs tested in normal colon, we observed that 940 maintained the same effect (in terms of size and direction) whereas 34 showed an opposite effect in tumor samples. Interestingly, of the 1,626 genes, 51 were cancer driver genes (CDG) (5 CRC specific; based on Cancer Gene Census [23]) but we did not find a significant enrichment of CDGs in triplets switching to causal compared to all other tumor triplets (**Fisher exact test p-value = 0.2903; odds-ratio = 1.274**). For 34 out of the 51 CDGs, we did not find a significant correlation between their expression and the expression of the corresponding TEs in normal samples pinpointing that these TEs have no impact on these genes in the normal state. Taken together, these results suggest an important role of TEs as drivers of gene expression during tumorigenesis.

### Non-coding germline variants activate driver TEs during tumorigenesis

We investigated whether any of the 9,714 tumor triplets were constituted of any previously identified tumor-specific or shared TE-eQTLs and assess how the model likelihood changed between normal and tumor. We identified 363 and 128 tumor triplets constituted of a shared or a tumor-specific TE-eQTL, respectively (**Figure 6A-B**) and observed that the 128 tumor triplets constituted with a tumor-specific TE-eQTL are significantly enriched for triplets switching to causal compared to the 363 tumor triplets constituted with a shared TE-eQTL (**Fisher Exact test p-value = 3.147e-05; Odds-ratio=2.4**) (**Figure 6B)**. Additionally, we observed that for 115 triplets with tumor-specific TE-eQTLs, the eQTL variant was not a significant eQTL for the corresponding gene in the triplet (**Figure 6C**), highlighting that the eQTLs get activated in the tumor state influencing TE expression that subsequently impact gene expression. Altogether, these results suggest that tumor-specific TE-eQTLs contribute to tumorigenesis by impacting genes through TEs, adding additional proof that germline variants can be contributing to tumorigenesis.

**Figure 6:**
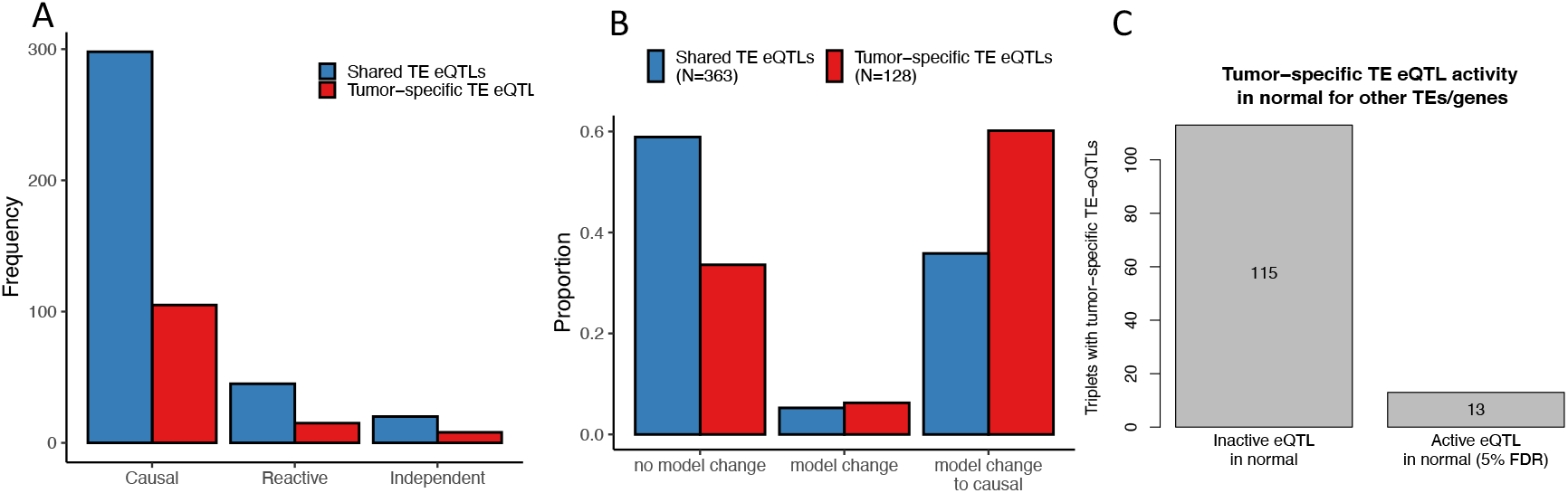
Tumor-specific and shared TE-eQTLs effects. **(A)** The barplot represent the frequency of the causal, reactive and independent model for the triplets with shared or tumor-specific TE-eQTLs (**B**) the barplot represents the model changes from normal to tumor for the triplets constituted of shared or tumor-specific TE-eQTLs. (**C**) represent the effect size of the shared and tumor-specific TE-eQTLs. (D) Barplot that represent the number of tumor-specific TE-eQTLs that are inactive eQTLs for the triplet associated gene.

### Driver TEs act as alternative promoters for genes in cancer

It has been shown that TEs could impact gene expression by acting as alternative promoters for nearby genes and creating chimeric transcripts (transpochimeric transcripts (tcGTs)) [24, 25]. To assess whether any of the tumor triplets with causal TEs were affected by tcGT events, we looked for cases where transcripts started from a TE and spliced into a single or multiple nearby genes (**methods**). We only kept tcGTs made up of the same TE and gene as in the 9,714 tumor triplets and that were significantly more abundant in tumor samples compared to normal samples using a Fisher exact test. At 5% FDR, we discovered 117 tcGTs present in 138 tumor triplets. Of these, 72 were causal and 46 switched to causal from normal to tumor. Interestingly, we detected tcGT events with a known tumor suppressor gene *RNF43* and two oncogenes *ETS2* and *SLCO1B3* supporting the extensive contribution of TEs during tumorigenesis.

## DISCUSSION

Transposable elements are important contributors to tumorigenesis and provide supplementary means by which gene expression can be altered in cancer. While many studies have used a hypothesis-driven approach and focused at specific TEs or their subfamilies for discovering TEs that alter the expression of nearby genes in cancer [26-28], applying a genome-wide scan could allow to obtain a better picture of the effects of TEs on gene expression during tumorigenesis.

Here, we present a global profile of tumor drivers and show that TEs are highly prevalent mediators of genetic effects on genes altering their expression, specifically in tumor. By combining genome and transcriptome data together we, show that TEs are under tight genetic control and discover that transcription factors regulate TE expression much more in tumor than in normal. By looking at the interplay between eQTL variants, transposable elements, and gene expression, we are able to dissect eQTL effects and show that for several genes, the genetic effect of an eQTL is passed on genes through TEs which act as mediators and drive gene expression. We observe this to occur significantly more in cancer than in normal and show that the majority of TEs increase the expression of affected nearby genes. Interestingly, we discover that TEs affecting known cancer driver genes in cancer have for most part no significant effect on these genes in normal suggesting a tumor-specific effect of these TEs. Additionally, in our study we show that alongside predisposing alleles and somatic mutations, germline variants are crucial contributors to tumorigenesis as these allow for transcriptional changes to occur at the level of TEs that in turn result in altered expression of nearby genes in cancer as shown previously [11].

While we focused on TEs impacting the expression of nearby genes in an independent manner, it is highly plausible that synergistic effects occur from both *cis-* and *trans*-acting TEs. Performing such an analysis could give a fuller picture of the regulatory network behind the regulation of gene expression through TE effects, requiring, however, a high sample size for sufficient statistical power. Nevertheless, because of the highly repetitive nature of transposable element sequences and their evolutionary relatedness among TE families, mapping short reads originating from TEs is a real challenge [18, 29]. Our RNA-seq dataset having a read length of 49bp, it is highly possible that we did not map all expressed TEs subsequently leading to missing information, as shown previously [18, 29]. Future studies where RNA-sequencing is performed with longer read lengths could allow for better mapping of expressed TEs and give us a fuller picture of the number of these driver TEs in cancer. Altogether, we have outlined that TEs are important mediators of genetic effects onto genes that could potentially be used as risk factors or new therapeutic targets for future drug development and aid in cancer treatment.

## METHODS

### 1.1. SYSCOL dataset

The Systems Biology of Colorectal cancer (SYSCOL) dataset contains data from genotypes and RNA-sequencing for matched normal-tumor samples (i.e., both tumor and normal samples originate from the same patient). Samples that had genotype data and molecular phenotype quantifications from tumor and normal (normal adjacent to tumor) tissue were analyzed, yielding 275 normal samples and 276 tumor samples. In case of multiple tumor samples from the same patient, only samples with quantifications from the most advanced tumor were kept.

### 1.2. Genotypes

We used imputed genotypes and only kept variants with a minor allele frequency (MAF) >=5%, yielding a total of 6,132,240 variants that were used for all downstream analyses.

### 1.3. Transcriptome quantifications

#### 1.3.1. Read mapping

SYSCOL samples were sequenced using 49bp, 75bp and 100bp read lengths. We first started by trimming all 75bp and 100bp reads down to 49bp to reduce any bias in downstream analysis stemming from read length. All trimmed samples were mapped to the human reference genome (hg37) using hisat2 [30]

#### 1.3.2. Transposable elements (TE) and genes quantifications

Gene and transposable element counts were generated using the *featureCounts* software [31]. We provided a single annotation file in *gtf* format to *featureCounts* containing both genes and transposable elements. This prevents any read assignation ambiguity to occur. For transposable elements, we used an in-house curated version of the *Repbase* database [12] where we merged fragmented LTR and internal segments belonging to a single integrant. We only used uniquely mapped reads for gene and TE counts. Molecular phenotypes that did not have at least one sample with 20 reads and for which the sum of reads across all samples was lower than the number of samples, were discarded. Furthermore, we normalized molecular phenotypes (TEs and genes) for sequencing depth using the TMM methodology as implemented in the *limma package* of Bioconductor [32] and used gene counts as library size for both TEs and genes. Finally, we removed any molecular phenotype that had more than 50% of missing data (zeros) in tumor and normal samples separately and took the union of molecular phenotypes, yielding 17’360 genes and 45’717 TEs for a total of 63’077 molecular phenotypes.

#### 1.3.3. Normalization of molecular phenotypes

The observed variability in molecular phenotypes from RNA-sequencing data can be of biological or technical origin. To correct for technical variability, while retaining biological variability, we residualised the molecular phenotype data for the covariates as described below:

1. To correct for population stratification that is observed between the SYSCOL samples, we used Principal Component analysis (PCA) results obtained from genotypes of SYSCOL patients. We only retained the first three principal components (PCs) as covariates.
2. In order to capture experimental/technical variability in the expression data, we performed PCA, centering and scaling, using *pca* mode from QTLtools software package [33]. To ascertain the number of PCs that capture technical variability, we used QTL mapping (see method 3.4.1 for the description of QTL mapping) to identify the best eQTL discovery power in both tumor and normal samples. To this end, we carried out multiple rounds of eQTL mapping for tumor and normal samples separately, each time using the 3 PCs from genotypes and incrementally adding 0, 1, 2, 5, 10, 20, 30, 40, 50, 60 and 70 PCs as covariates. This approach resulted in identifying 30 PCs in tumor and normal samples for maximizing eQTL discovery.

In total, 33 covariates were regressed out from tumor and normal sample expression data using QTLtools *correct* mode [33]. We additionally rank-normalized on a per phenotype basis across all samples such that quantifications followed normal distribution with mean 0 and standard deviation 1 N(0,1) using QTLtools *--normal* option [33].

### 1.4. DNA Methylation data and differential methylation of eQTLs

We used microarray based DNA methylation data from the SYSCOL project and a similar approach to a previous study to find differential methylation of eQTLs [11]. In brief, we calculated the absolute value difference of the medians of normalized methylation probe betas in normal and tumor that we call median differential methylation. We then compared the distribution of there medians in tumor-specific TE and genes eQTLs vs. the shared TE and gene eQTLs and calculated a P-value using the Mann Whitney U test. P-values were corrected for multiple testing using the *R/qvalue* package with a given FDR threshold of 5%.

### 1.5. Differential TE/gene expression analysis

The DESeq2 R package [34] was used in calculating differentially expressed genes and TEs. We normalized the raw TE/gene counts within the DESeq2 package using the sequencing date, GC mean and insert size as covariates. The differential expression P-values were corrected for multiple testing using an FDR threshold of 5%.

### 1.6. Transcriptome QTL analysis

All analyses were performed separately for normal and tumor samples. We used imputed genotypes with MAF >=5 %, gene expression data with normalized counts per million (CPMs) (as described above) for both eQTL and conditional eQTL mapping.

#### 1.6.1. Expression Quantitative Trait Loci (eQTL) mapping

For eQTL mapping, we used *cis* mode of the QTLtools software package [33]. For each molecular phenotype:

1. We counted all genetic variants in a 1 Mb window (+/- 1 Mb) around the transcription start site (TSS) of the phenotype and tested all variants within this window for association with the phenotype. We only retained the best hits which are defined as the ones with the smallest nominal p-value.
2. Next, we used permutations to adjust the nominal p-values for the number of variants tested. More specifically, we randomly shuffled the quantifications of the phenotypes 1’000 times and retained only the most significant associations. This created a null distribution of 1’000 null p-values. Then, we fitted a beta distribution on the null distribution and used the resulting beta distribution to adjust the nominal p-value. Principally, this strategy allows to quantify the chance of getting a smaller p-value than the nominal one in random datasets.

This effectively gave us the best variant in *cis* together with the corresponding adjusted p-value of association for each molecular phenotype. Finally, to correct for the number of phenotypes being tested we used False Discovery Rate (FDR) correction approach. More specifically, we used the *R/qvalue* package [35] to perform genome-wide FDR correction which ultimately facilitated to extract all phenotype-variant pairs that are significant at a pre-determined FDR threshold, 5% FDR in this case.

#### 1.6.2. Conditional analysis for eQTL mapping

The *cis* mode informs us on the best phenotype-variant pair only. Given that the expression of molecular phenotypes can be affected by multiple *cis* eQTLs, we proceeded with conditional analysis to discover all eQTLs with independent functional effects on a given phenotype. Principally, new discoveries are made after conditioning on previous ones. Again, *cis* mode in the QTLtools software package was used [33]. In brief, after running permutations (**method 1.4.1**) for each phenotype, we calculated a nominal p-value threshold of being significant. We first determined the adjusted p-value threshold that corresponds to the targeted FDR level and then used the beta quantile function to go from adjusted p-value to a specific nominal p-value threshold. For conditional analysis, forward-backward methodology is used to discover all independent QTLs and to identify the most likely candidate variants, while at the same time controlling for a given FDR (5% FDR in this case). We only kept the top variant for each signal.

#### 1.6.3. Tissue-specific and shared eQTL analysis

To discover tissue specific and shared eQTLs, we used the eQTL results obtained after running the conditional pass. In total, we tested 17’077 eQTLs to discover normal-specific eQTLs and 6’591 to discover tumor-specific eQTLs. To do that, we used linear mix models using an interaction term between dosage and tissue (i.e tumor or normal) to test whether the slopes in normal and tumor are significantly different. Linear mix models are needed here because normal and tumor samples are originating from the same patient thus genotypes will be identical. We did this for tumor and normal eQTLs separately. Then we performed multiple test correction using the *R/qvalue package* [35] with a given FDR threshold of 5%. Additionally, for all significant results at 5% FDR, if eQTL slopes (slopes given from conditional QTL mapping using *QTLtools*) in normal and tumor had the same direction, then we only kept the ones where the SNP-phenotype association in the opposite tissue was not nominally significant (P>0.05) as given by the *cis nominal pass* mode in the QTLtools package [33]. Shared eQTLs are defined as the ones where the P-value for the interaction term is not significant but need to be significant eQTLs (5% FDR) in both normal and tumor as assessed by the conditional QTL mapping.

#### 1.6.4. Functional enrichment analysis

To compare the QTL variants to a null distribution of similar variants without regulatory association, we sampled for each eQTL variant 100 random regulatory genetic variants matching for relative distance to TSS (withing 2.5kb) and minor allele frequency (within 2%) and only kept variants that are not eQTLs for any other TE or gene (nominal p-value > 0.05). The enrichment for a given category was calculated as the proportion between the number of regulatory associations in a given category and all regulatory variants over the same proportion in the null distribution of variants. The p-value for this enrichment is calculated with the Fisher exact test. Finally, we corrected for multiple testing using an FDR threshold of 5% using the “*p*.*adjust*” function in the R programming language.

We used two different datasets for the functional enrichment. For gene and TE eQTLs in normal and tumor, we used available transcription factor (TF) ChIP-seq data from Ensembl Regulatory Build [15]. For each TF, we combined all peaks from all available cell-types. Regarding the tumor-specific vs. shared TE and gene eQTLs, we used available ChIP-seq data from the colorectal cancer LoVo cell line [19].

##### Testing for associations between TEs and genes

To discover associations between TEs and genes, we proceeded in a similar way to what we did for QTL mapping (**method 1.4.1**). Effectively, we used TE expression as our “genotypes” and genes as our phenotype. Then, we corrected for multiple testing using the *R/qvalue* package with a given FDR of 1%. We then estimated the nominal p-value thresholds for each phenotype being tested as described in (method 1.4.2) with a given FDR of 1%. Given the nominal threshold we get for each gene, we then extracted all TEs with an association P-value below this threshold which could give multiple TEs for a gene in some cases.

### 1.7. Quantifying TE-gene pairs

To quantify each of TE-gene pairs that have been found to be significant, we used a dimensionality reduction approach based on PCA as previously described [22]. Specifically, for each TE-gene pair, we aggregated gene expression together with TE expression by using the coordinates on the first PC. This effectively built a quantification matrix with rows and columns corresponding to the number of TE-Gene pairs and individuals, respectively. All quantifications have been rank-normalized on a per phenotype basis so that the values match a normal distribution N(0,1). This prevents outlier effects in downstream association testing. This is all implemented in the *clomics* software package [22].TE

### 1.8. Causal inference by Bayesian networks for QTL-TE-Gene triplets

Bayesian networks (BNs) are a type of probabilistic graphical model that uses Bayesian inference to compute probabilities. BNs aim to model conditional dependencies and therefore causation by representing conditional dependencies as edges and random variables as nodes in a directed acyclic graph. The flow of information between two nodes is reflected by the direction of the edges, giving an idea of their causal relationship. BNs have been previously used in a genetic framework [20] to get insight into the most likely network from which the observed data originates.

In BNs, the joint probability density can be divided into marginal probability functions and conditional probability functions for the nodes and edges, respectively. Additionally, BNs satisfy the local Markov property where each variable is conditionally independent of its non-descendants given its parent variables. In the context of this study, we used BNs to learn the causal relationships between triplets of variables, each one containing a genetic variant, a transposable element and a gene. In practice, only three distinct network topologies where relevant to the hypotheses we wanted to test (**Supplementary figure 12**). More specifically, we looked at:

1. The **causal** scenario where the genetic variant affects first the TE and then the gene.
2. The **reactive** scenario where the genetic variant affects the gene first and then the TE.
3. The **independent** scenario in which the variant affects the gene and the TE independently.

Of note, we only retained network topologies that assume that the signal systematically originates from the genetic variant. In practice, we applied BNs on data that was obtained from running an QTL mapping using the TE-gene pairs using a similar approach to QTL mapping described above (**Method 1.4.1**) and only kept significant results at 5% FDR which corresponds to 11’425 QTL-TE-gene triplets in normal and 9’488 QTL-TE-gene triplets in tumor.

For each triplet, we build a 275 × 3 matrix in normal and 276 × 3 matrix in tumor containing normalized quantifications and used it to compute the likelihood of the 3 BN topologies using the *R/bnlearn* package (*Version 4*.*5*) [36]. As a last step, we went from likelihoods to posterior probabilities by assuming a uniform prior probability on the three possible topologies. Posterior probabilities where used for all BN-related analyses.

### 1.9. Transpochimeric transcripts analysis

First, a per sample transcriptome was computed from the RNA-seq bam file using StringTie [37] with parameters –j 1 –c 1. Each transcriptome was then crossed using BEDTools [38] to both the ensembl hg19 coding exons and curated RepBase [12] to extract TcGTs for each sample. Second, a custom python program was used to annotate and aggregate the sample level TcGTs into counts per groups (normal, tumor). In brief, for each dataset, a GTF containing all annotated TcGTs was created and TcGTs having their first exon overlapping an annotated gene or TSS not overlapping a TE were discarded. From this filtered file, TcGTs associated with the same gene and having a TSS 100bp within each other were aggregated. Finally, for each aggregate, its occurrence per group was computed.

## AUTHOR CONTRIBUTIONS

N.M.R.L, H.O and E.T.D designed the study. N.M.R.L analyzed the data and wrote the manuscript and N.M.R.L, H.O, D.T and E.T.D interpreted the results. E.P shared the quantifications data.

## COMPETING INTERESTS

Emmanouil T. Dermitzakis is currently an employee of GSK. The work presented in this manuscript was performed before he joined GSK. All other authors declare no competing interests.

## ACKNOWLEDGMENTS

The computations were performed at the University of Geneva on the Baobab cluster. This work was supported by grants from Louis-Jeantet Foundation support (to E.T.D) and SNSF grant (to E.T.D). The funders had no role in study design, data collection and analysis, decision to publish, or preparation of the manuscript.

