## Supplementary Figures for "Transposable elements mediate genetic effects altering the expression of nearby genes in colorectal cancer"

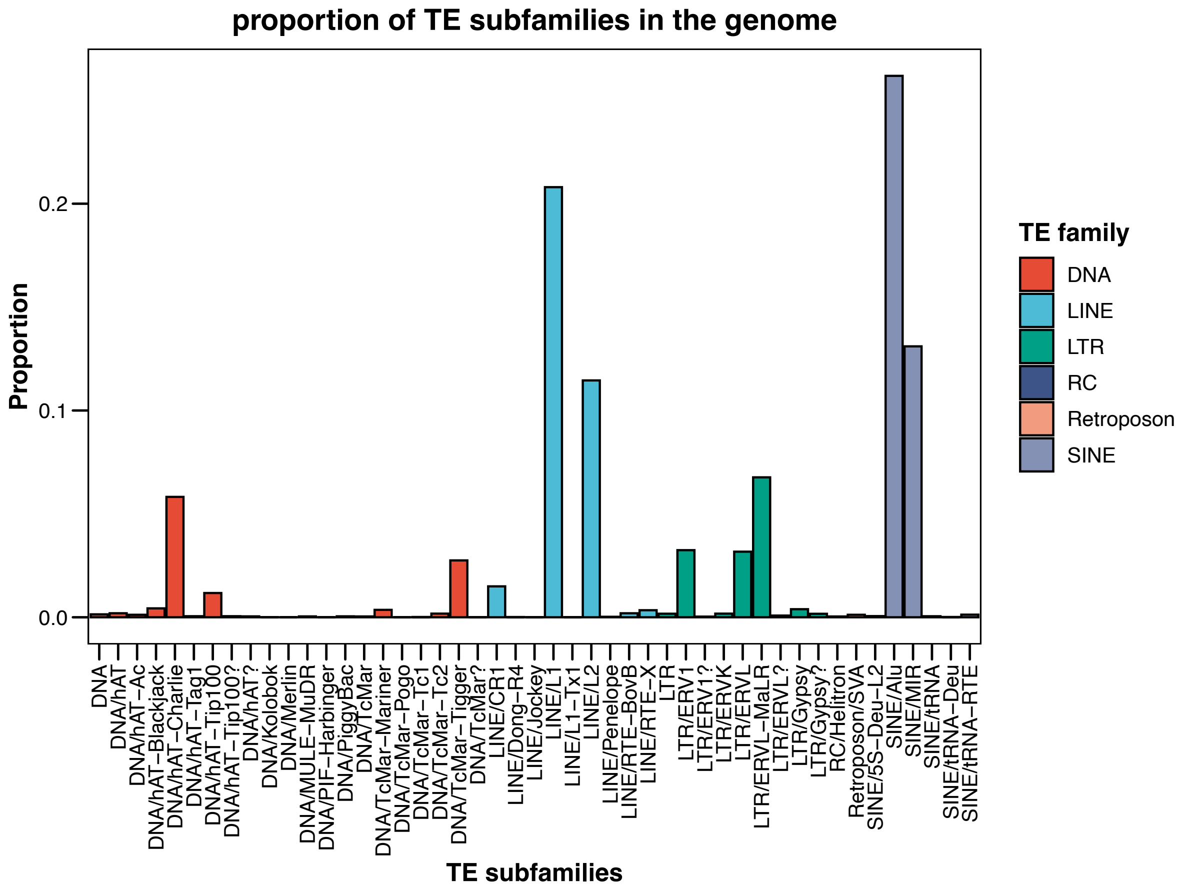


**Supplementary figure 1 |** ***Proportion of TE subfamilies in the human genome.*** The majority of transposable elements in the human genome are Alu and tRNA from the SINE family, followed by L1 and L2 elements from the LINE family and ERV1, ERVL and ERVL-MaLR from the LTR family.

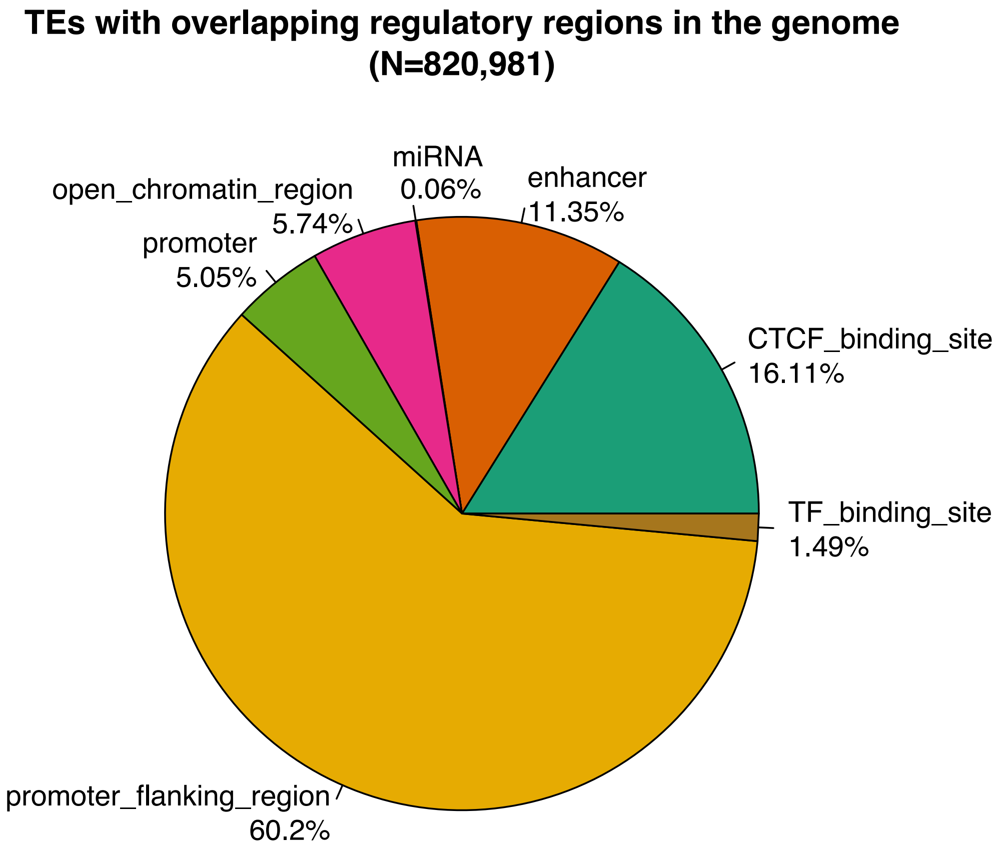


**Supplementary Figure 2 | TEs overlapping regulatory regions in the human genome**. Pie plot representing the proportion of transposable elements overlapping with each regulatory regions. Of the ~4.6 million TEs in the human genome, 820,981 TEs are overlapping with at least one regulatory region.

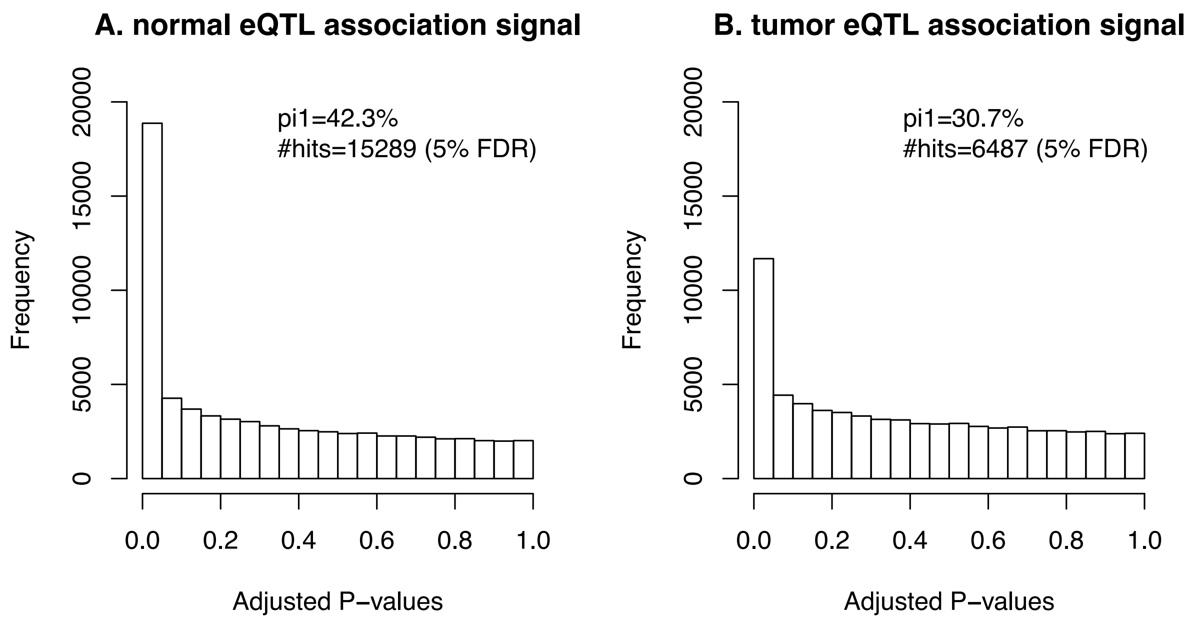


**Supplementary figure 3 | *cis eQTL discovery p-value distribution*.** Histograms of p-value distribution of *cis*- eQTL discovery in (**A**) normal and (**B**) tumor.

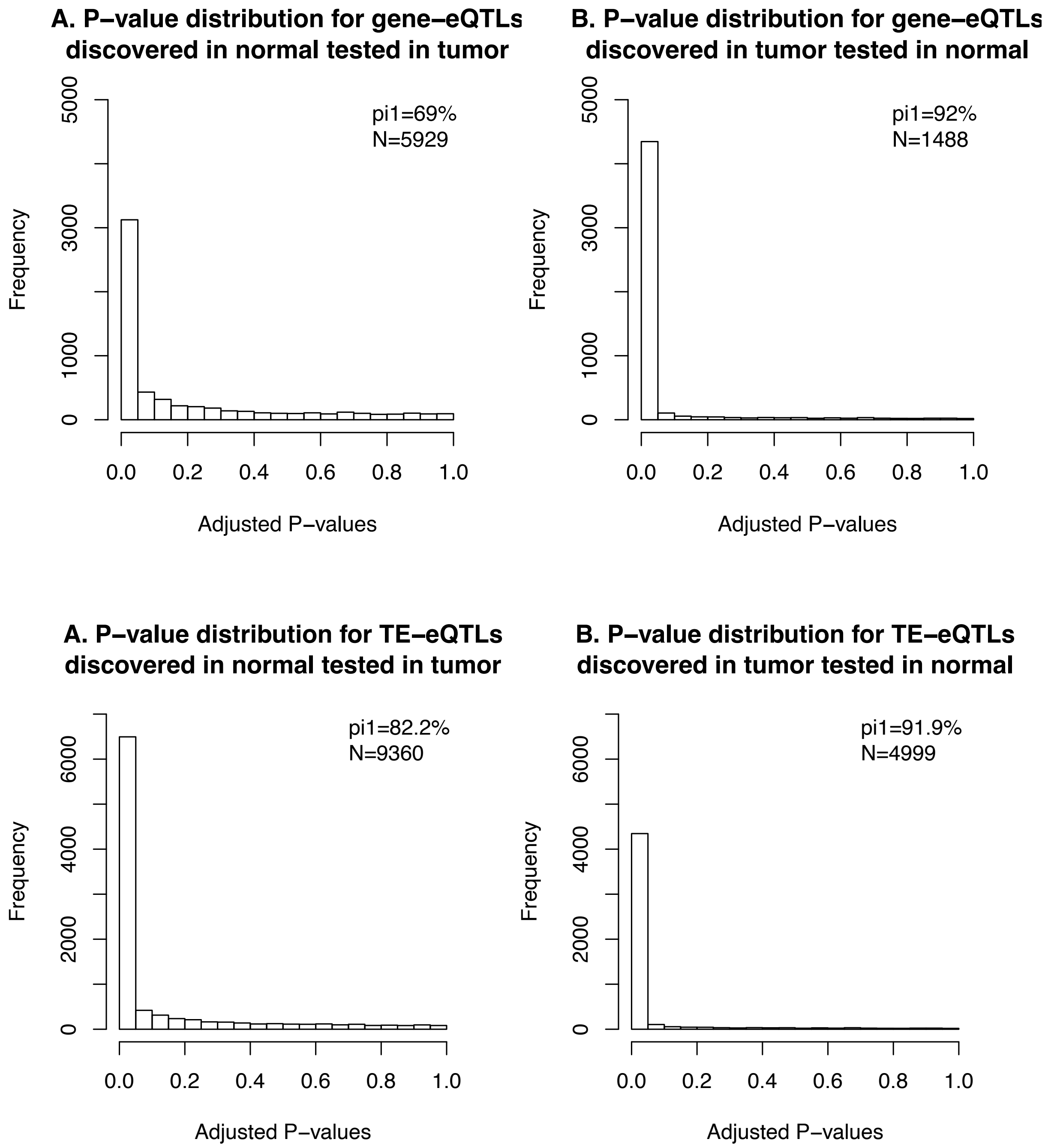


**Supplementary figure 4 | P-value distributions of significant SNP-gene or SNP-TE pairs tested in the other tissue.** The $\pi_1$ statistic estimates the tissue sharing of eQTLs. We observe that SNP-TE and SNP-gene pairs discovered in tumor are replicated better in normal than normal eQTLs tested in tumor.

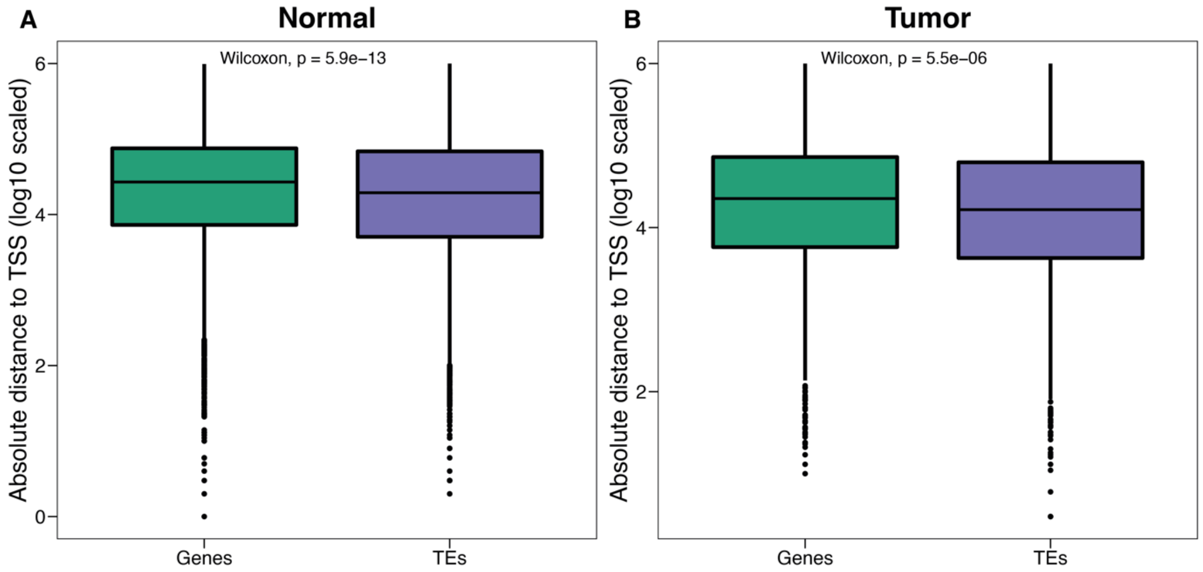


**Supplementary figure 5 | TE and gene eQTL distance to TSS in normal and tumor.** We observe that in both normal and tumor, TE-eQTLs are closer to the TSS of TEs than gene-eQTLs are to the TSS of genes. This could be because of the smaller evolutionary time TEs have in the human genome compared to genes, making local effects much more likely to occur.

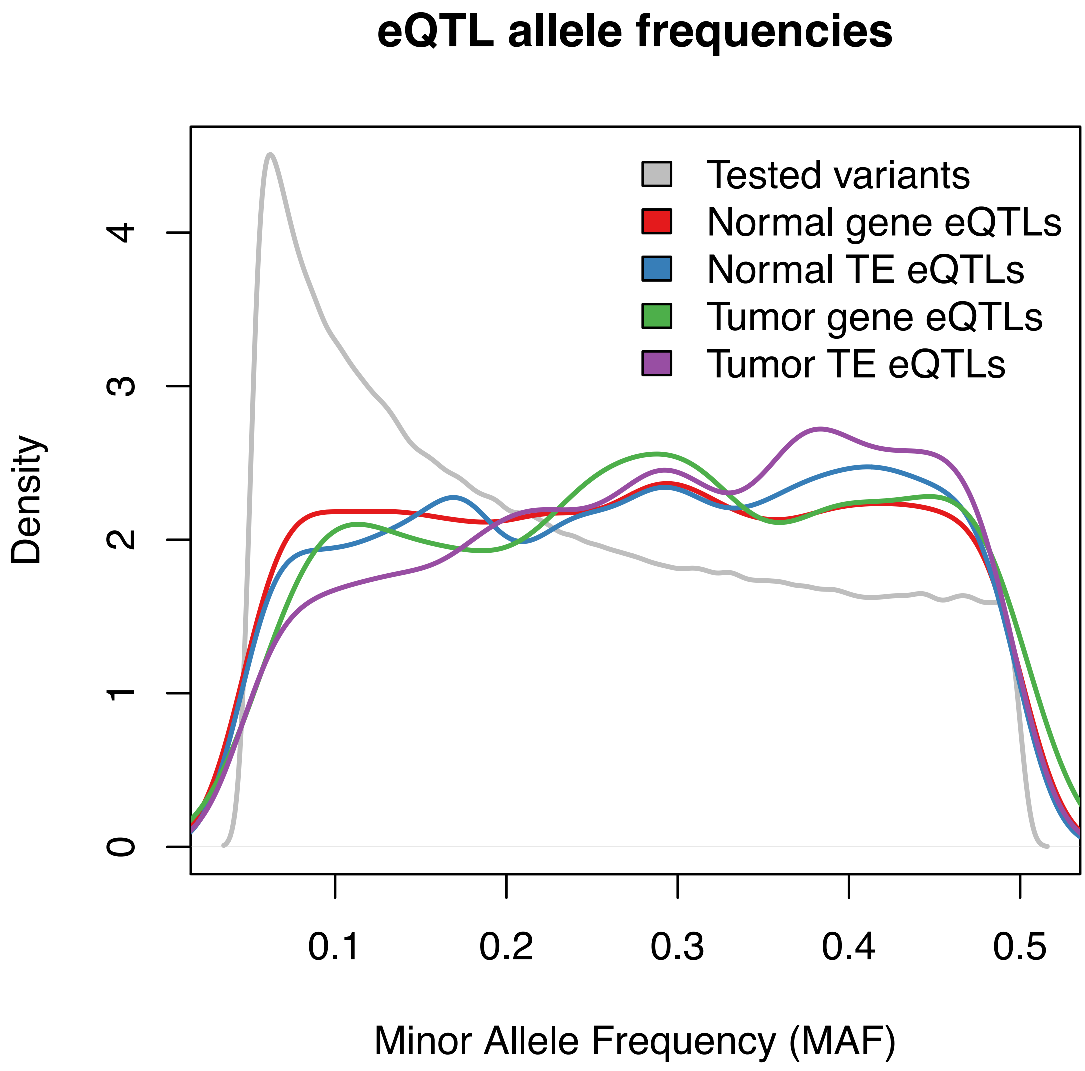


**Supplementary figure 6 | eQTL variant allele frequencies.** We observe that the allele frequencies for the TE- and gene-eQTLs are very similar in normal and tumor.

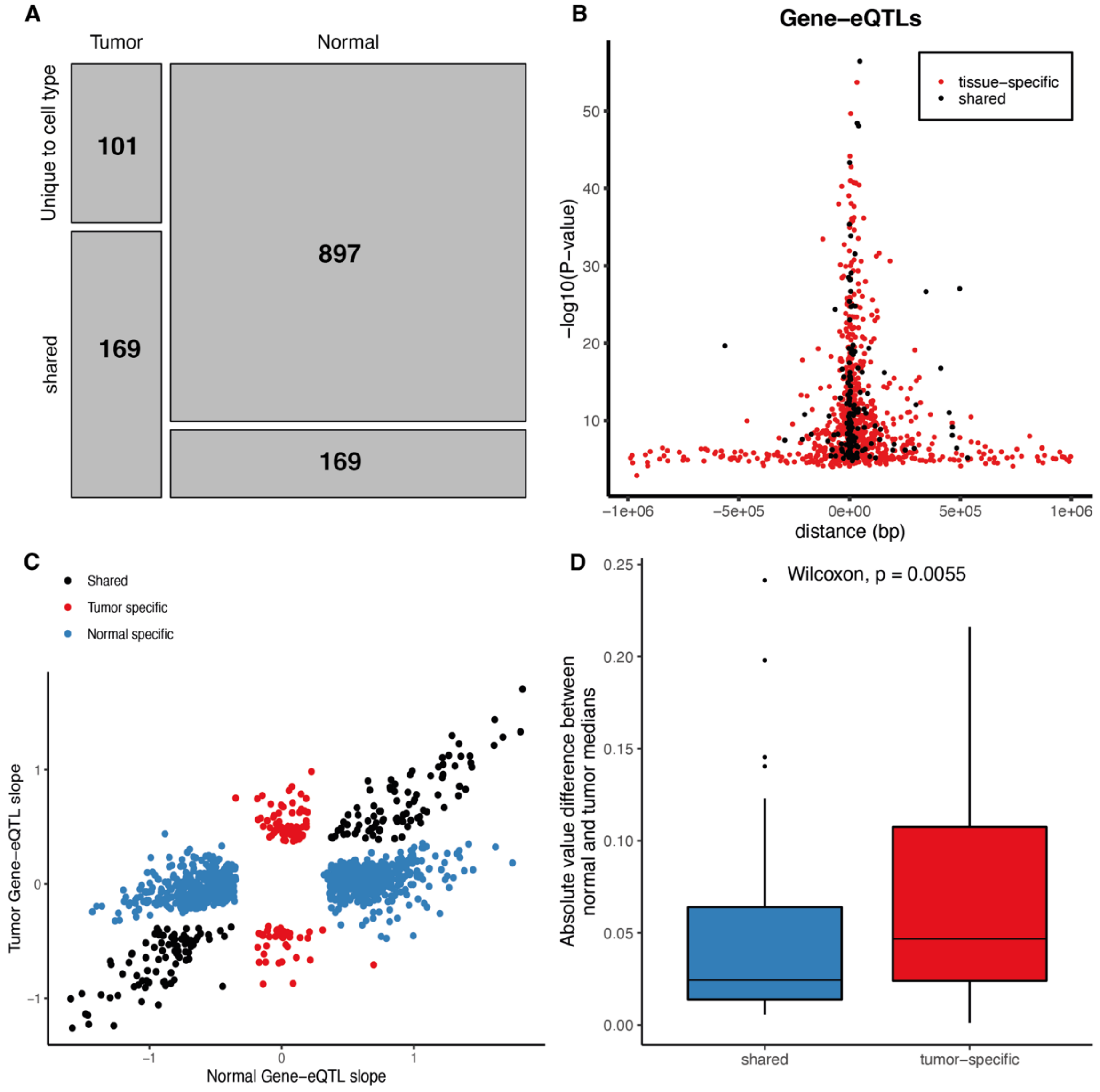


**Supplementary figure 7 | Tissue specificity of Gene-eQTLs**. (**A**) Mosaic plot of tissue specificity of Gene-eQTLs. (**B**) Tissue specificity and distance of Gene-eQTL to transcription start site (TSS). The shared Gene-eQTLs (black) are closer to the TSS than are the tissue specific Gene-eQTLs (red) (Wilcoxon P<2.2e-16). (**C**) Gene-eQTL slopes for the normal specific Gene-eQTLs in blue, the tumor specific in red and shared in black. (**D**) Boxplot of the absolute value difference of median methylation betas between normal and tumor samples for shared and tumor-specific gene-eQTLs.

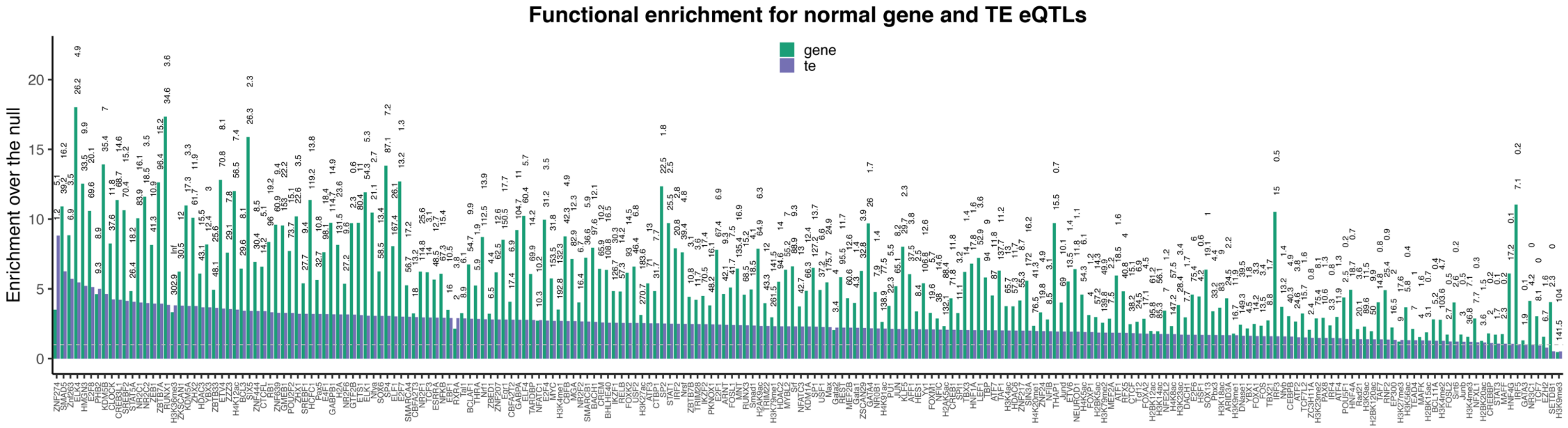


**Supplementary Figure 8 | Functional enrichment for gene- and TE-eQTLs in normal**. Enrichment over the null for gene- and TE-eQTLs in normal. Only plotting cases where the enrichment was significant at 5% FDR for either gene- or TE-eQTLs.

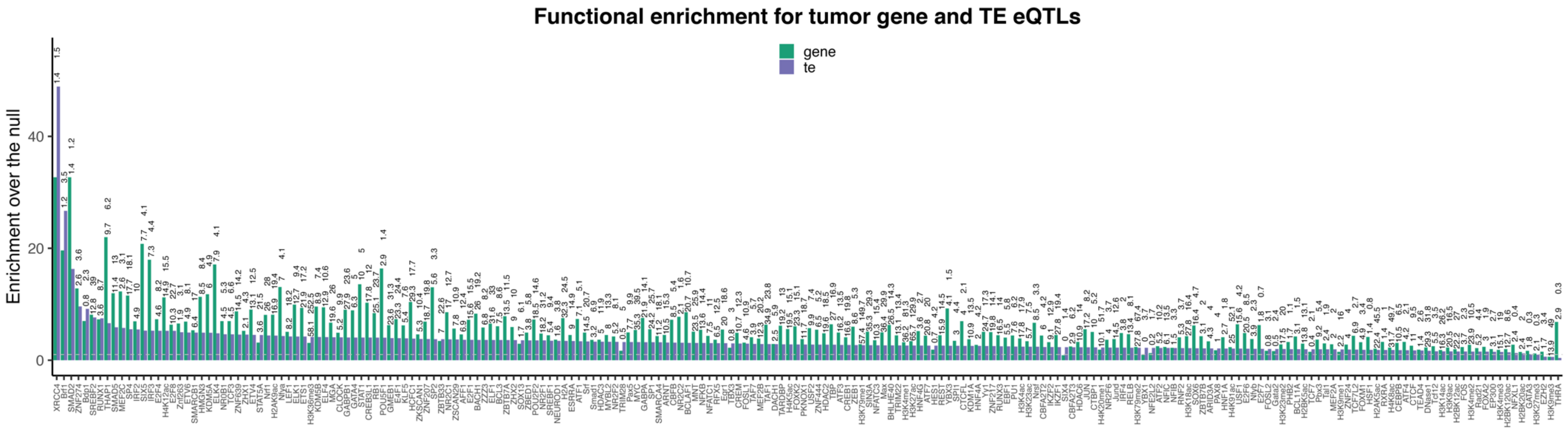


**Supplementary Figure 9 | Functional enrichment for gene- and TE-eQTLs in tumor**. Enrichment over the null for gene- and TE-eQTLs in tumor. Only plotting cases where the enrichment was significant at 5% FDR for either gene- or TE-eQTLs.

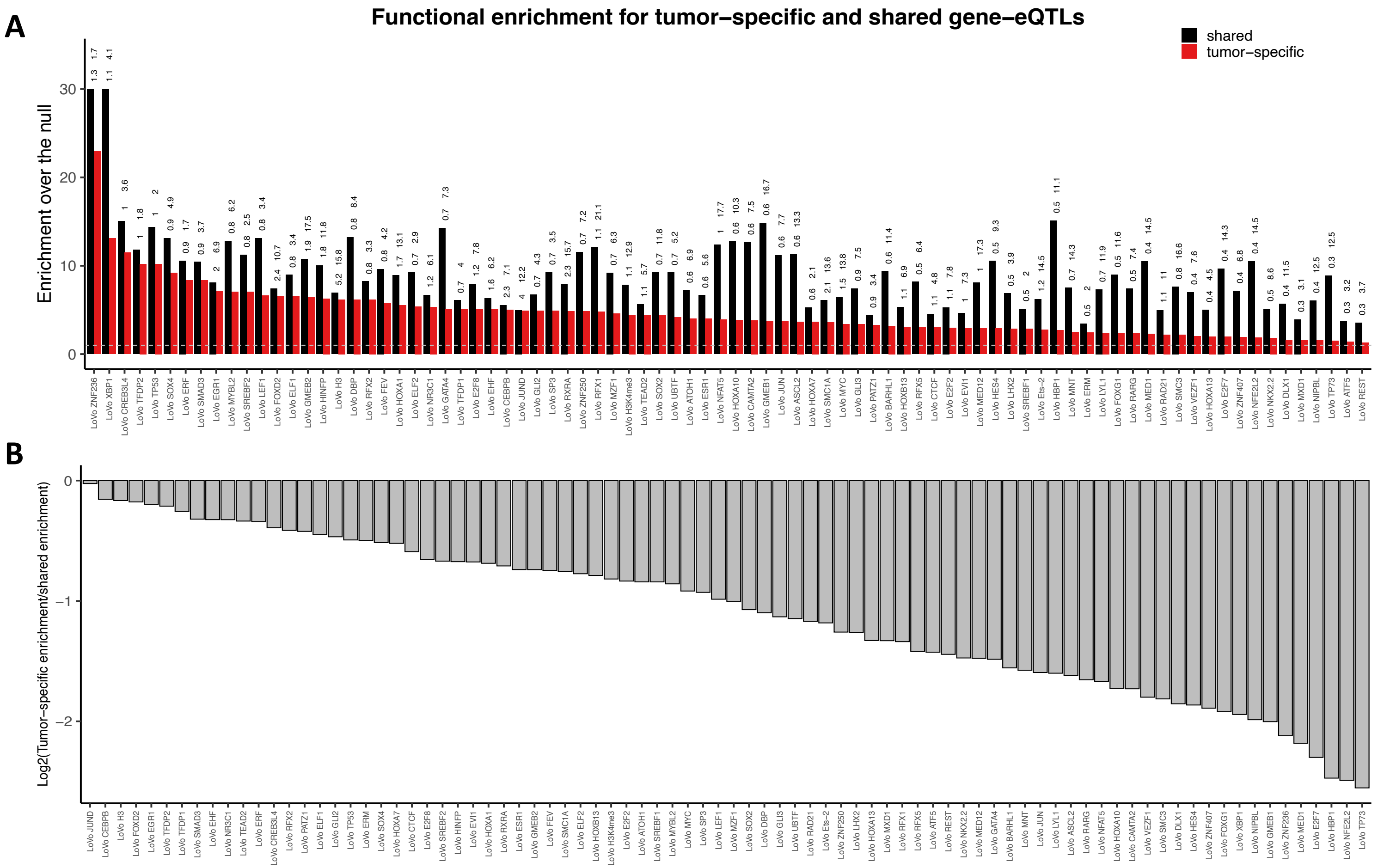


**Supplementary figure 10 | Functional enrichment for Tumor-specific and shared gene-eQTLs.** (A) Enrichment over the null for tumor-specific and shared gene-eQTLs. (B) log2 ratio between tumor-specific enrichment and shared enrichment. Only plotting cases where the enrichment was significant at 5% FDR for either tumor-specific or shared gene-eQTLs. None of the TFs have a stronger enrichment for tumor-specific gene-eQTLs indicating that these TFs regulate gene expression in both the normal and tumor state.

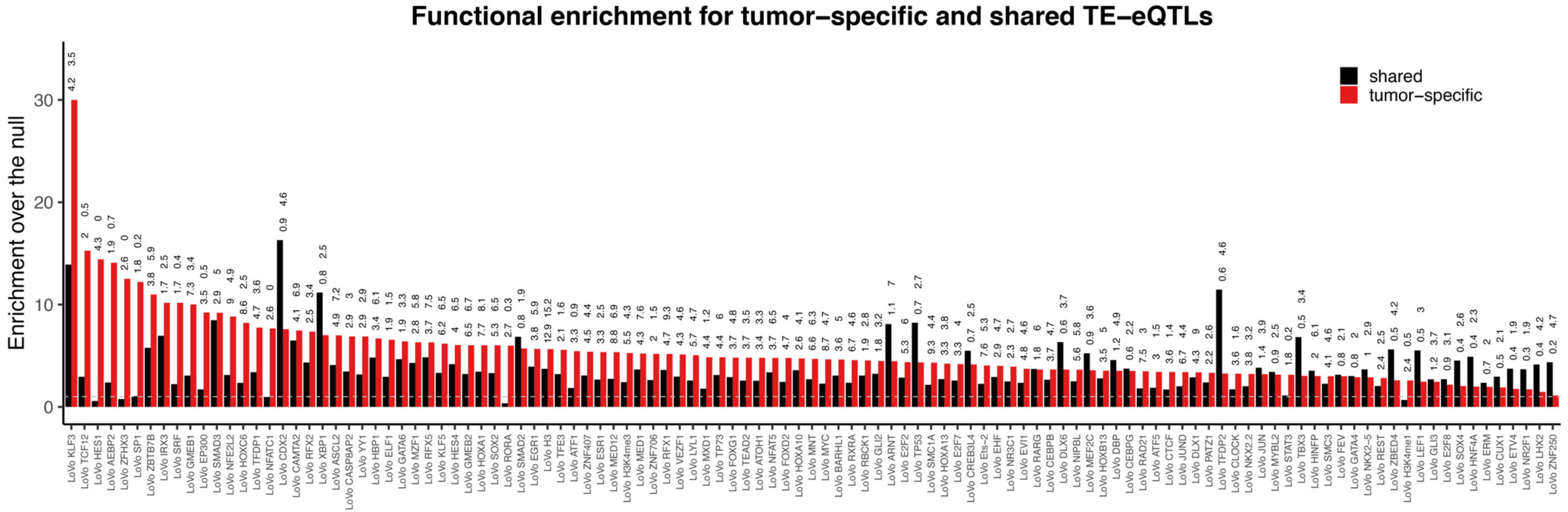


**Supplementary figure 11 | Functional enrichment for Tumor-specific and shared TE-eQTLs.** (A) Enrichment over the null for tumor-specific and shared TE-eQTLs. Only plotting cases where the enrichment was significant at 5% FDR for either tumor-specific or shared TE-eQTLs.

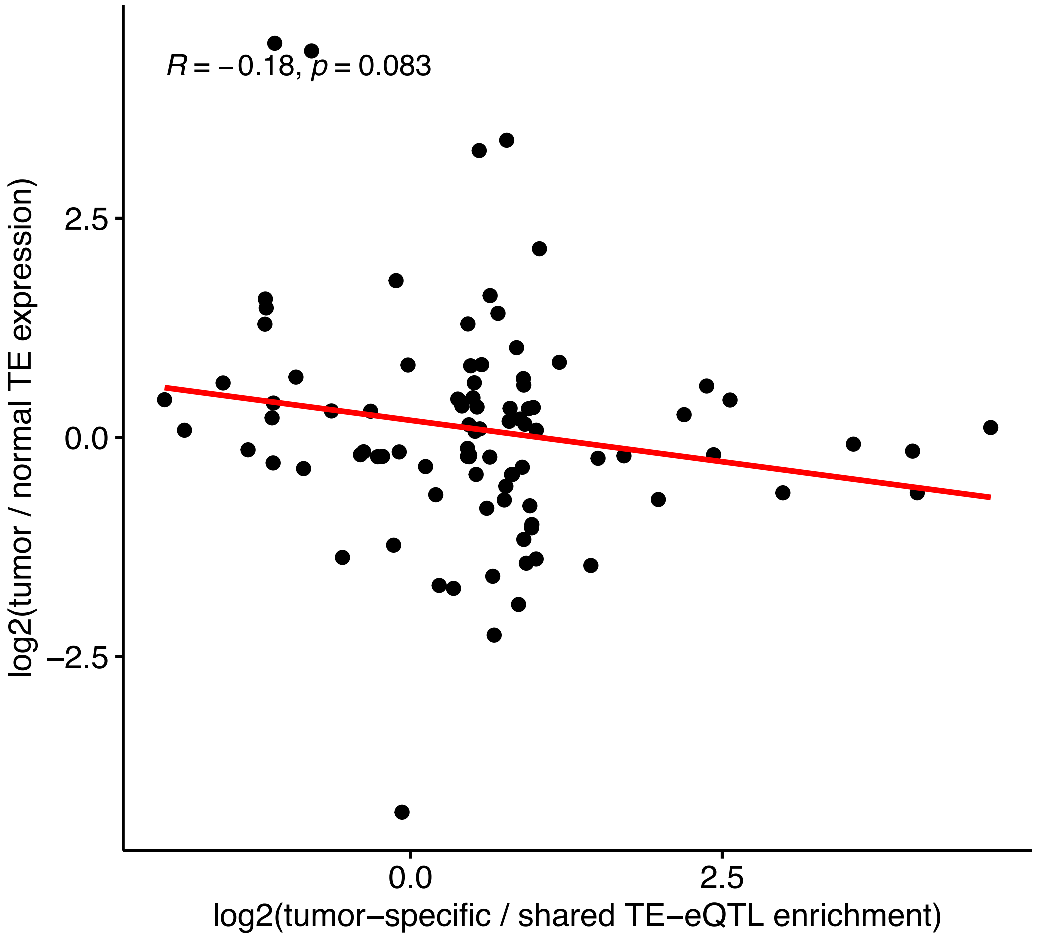


**Supplementary figure 12 | Correlation of differential enrichment of functional binding sites and differential expression of the corresponding transcription factors.** We compared the ratio of tumor expression over the normal expression of differentially expressed transcription factors to the ratio of enrichment for the binding sites of the same transcription factors in tumor specific TE-eQTLs over the shared TE-eQTLs. We find no significant correlation indicating that differential expression of the corresponding TFs do not drive the tumor-specific TE-eQTLs.

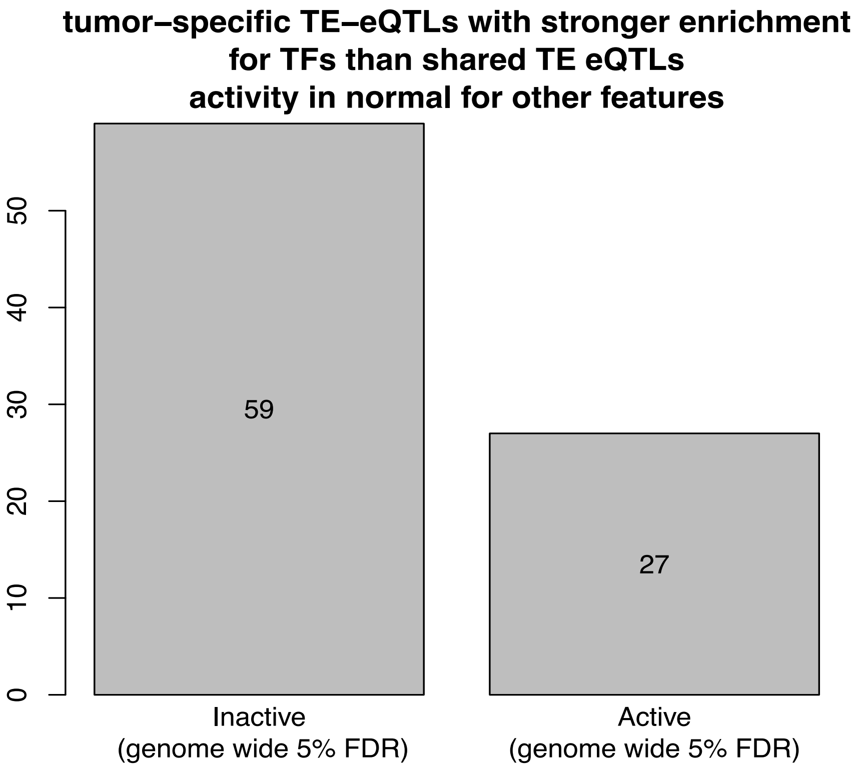


**Supplementary figure 13 | Tumor-specific TE-eQTLs overlapping with the enriched transcription factors and their activity in normal regarding other genes/TEs.** We checked whether any of the tumor-specific TE-eQTLs overlapping with the transcription factors we found to have a stronger enrichment for tumor-specific TE-eQTLs are active or inactive eQTLs for other TEs or genes in *cis*. We discovered that 59 of them are not significant eQTLs for any TE or gene in normal indicating that these regions are probably inactive and get activated in tumorigenesis.

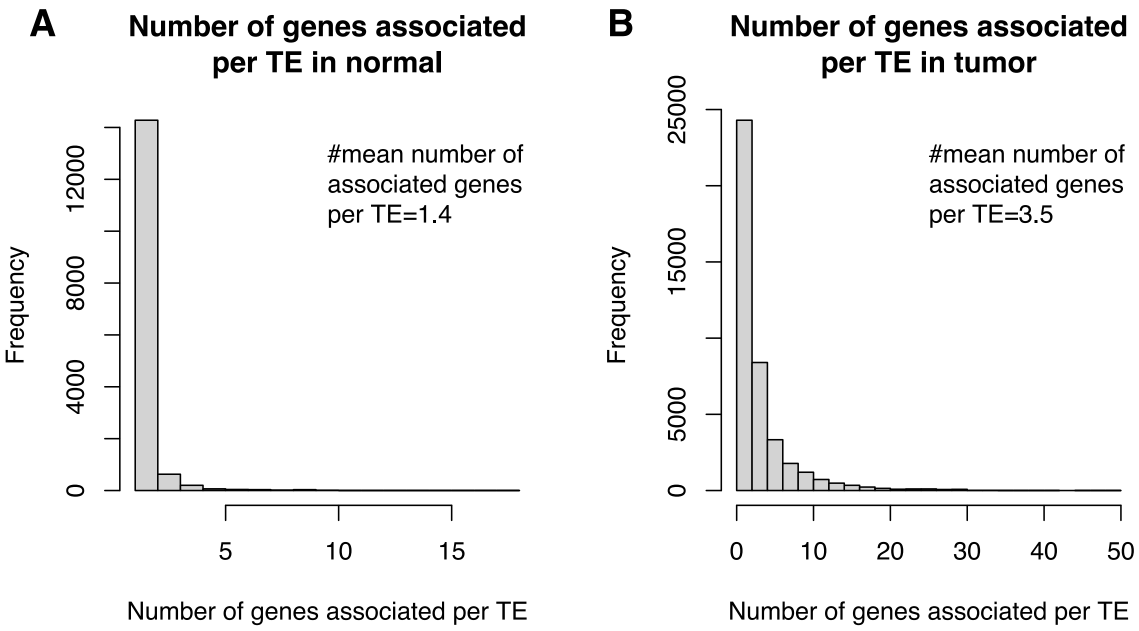


**Supplementary figure 14 | Mean number of genes associated per TE in (A) normal and (B) tumor.**

We observe that in normal there are less genes associated per TE (mean number of associated genes per TE = 1.4) compared to tumor (mean number of associated genes per TE = 3.5)

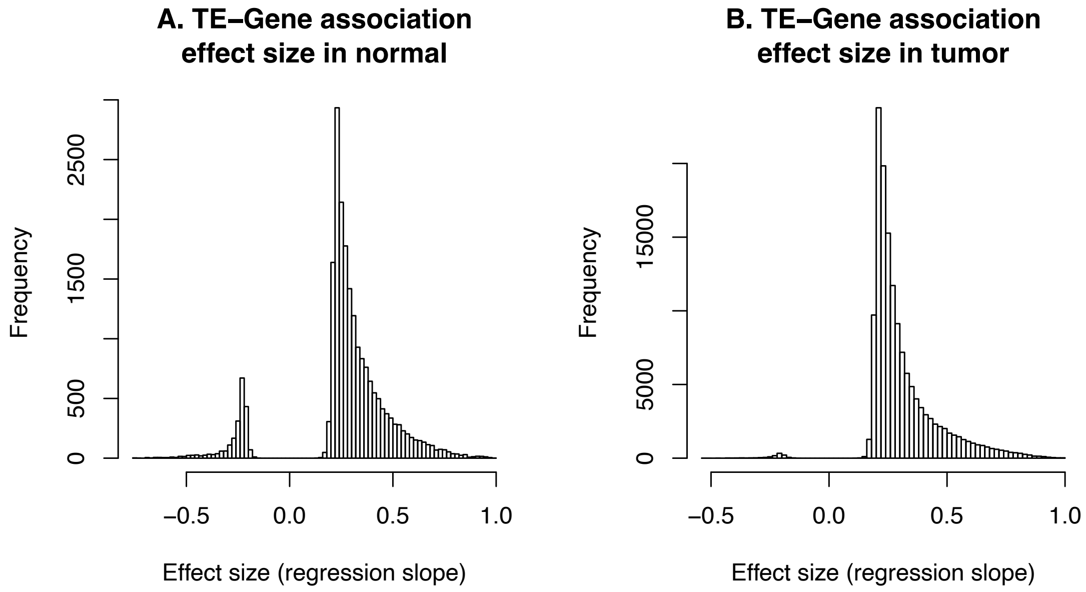


**Supplementary figure 15 | TE-gene effect sizes (regression slope) in (A) normal and (B) tumor.**

We observe that most TEs are positively associated with a gene.

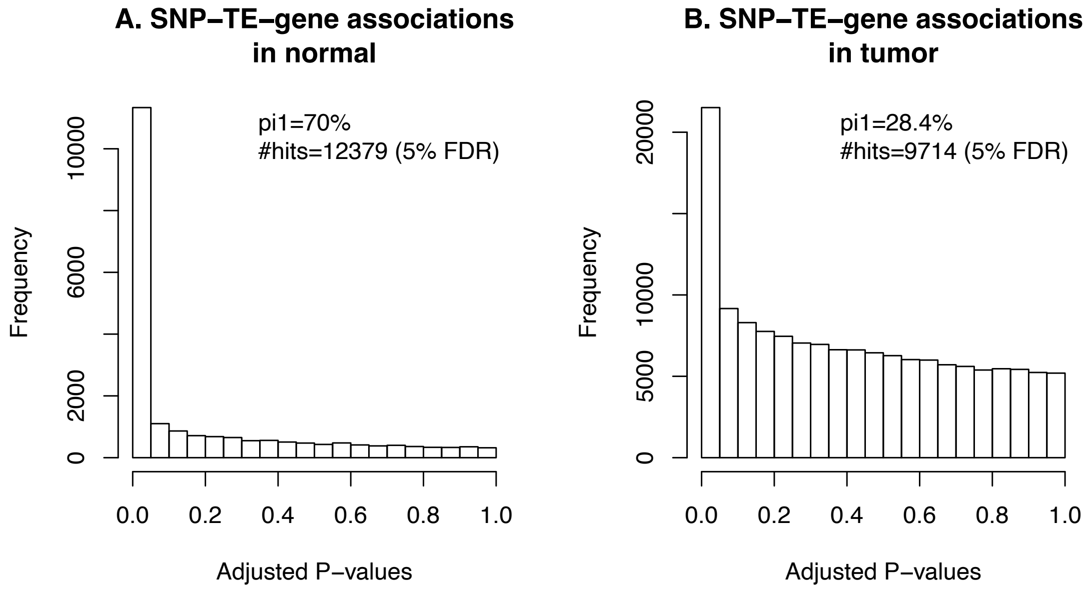


**Supplementary figure 16 | P-value distribution of eQTL TE-gene discovery in (A) normal and (B) tumor.** We observe that in normal (A) we discovered 12,379 triplets and in (B) tumor 9,714 triplets at 5% FDR.

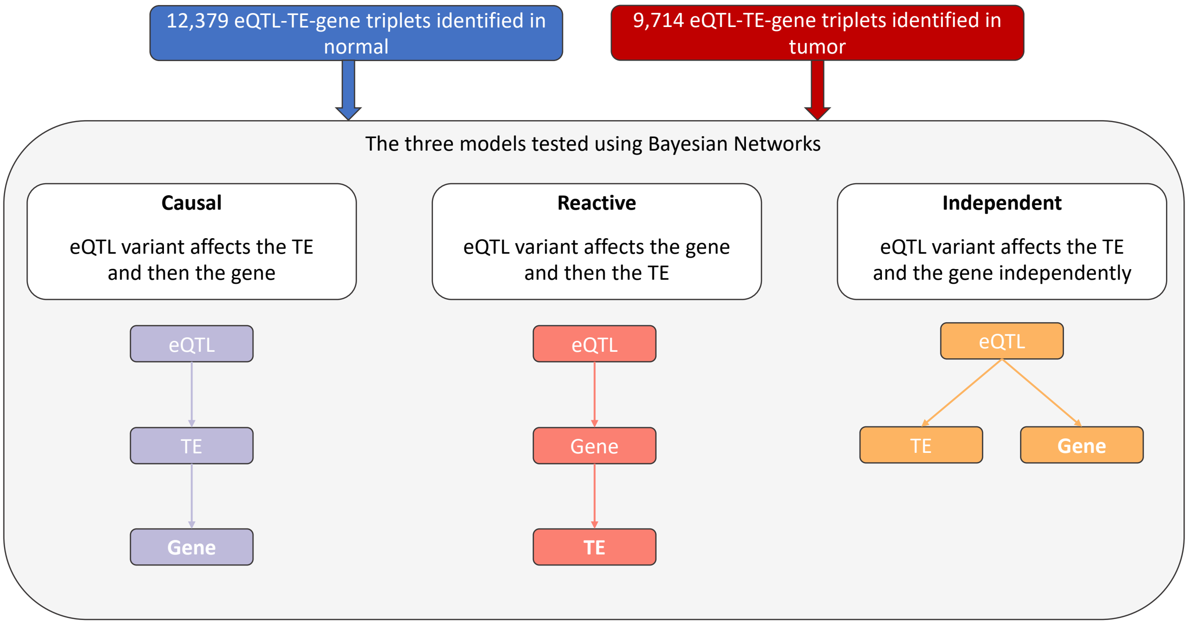


**Supplementary figure 17 | Causal relationship of eQTLs, TEs and genes approach**. To infer the most likely causal relationship between eQTL variants, TEs and genes, we tested three models using Bayesian Networks (BNs). The causal model where the eQTL variant affects the TE and then the gene, the Reactive model where the eQTL variants affects the gene and then the TE and the Independent model where the eQTL variant affects the TE and gene independently. For each triplet we obtained log likelihoods and we calculated posterior probabilities using a uniform prior probability for each of the three models.

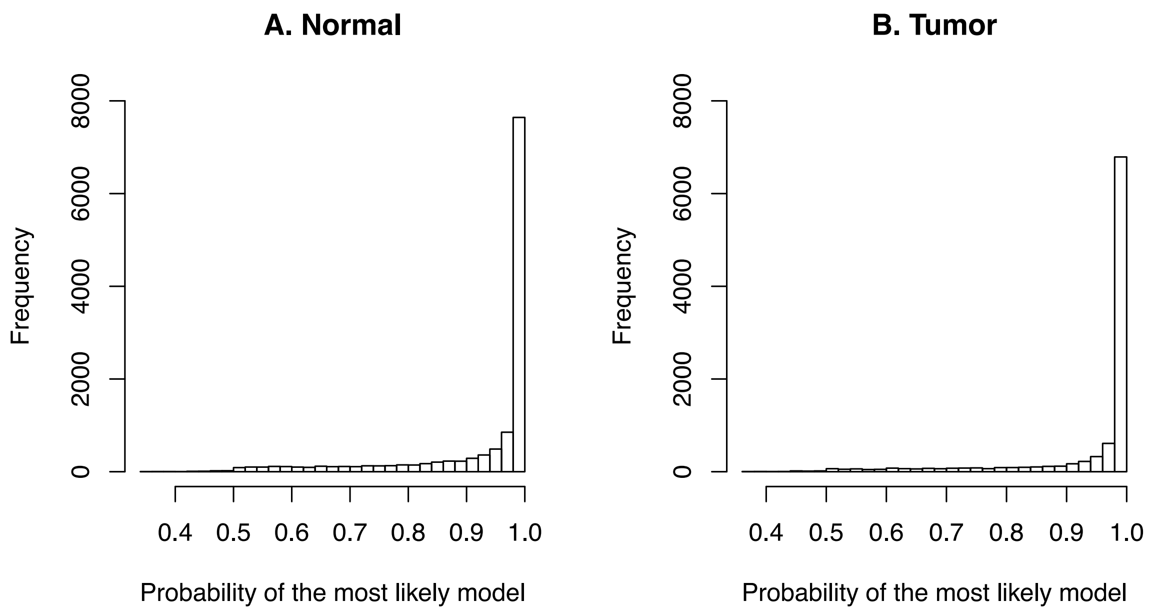


**Supplementary figure 18 | Probability of the most likely model in (A) normal and (B) tumor.** We can see that the probabilities of the most likely models are for most part above 0.8.

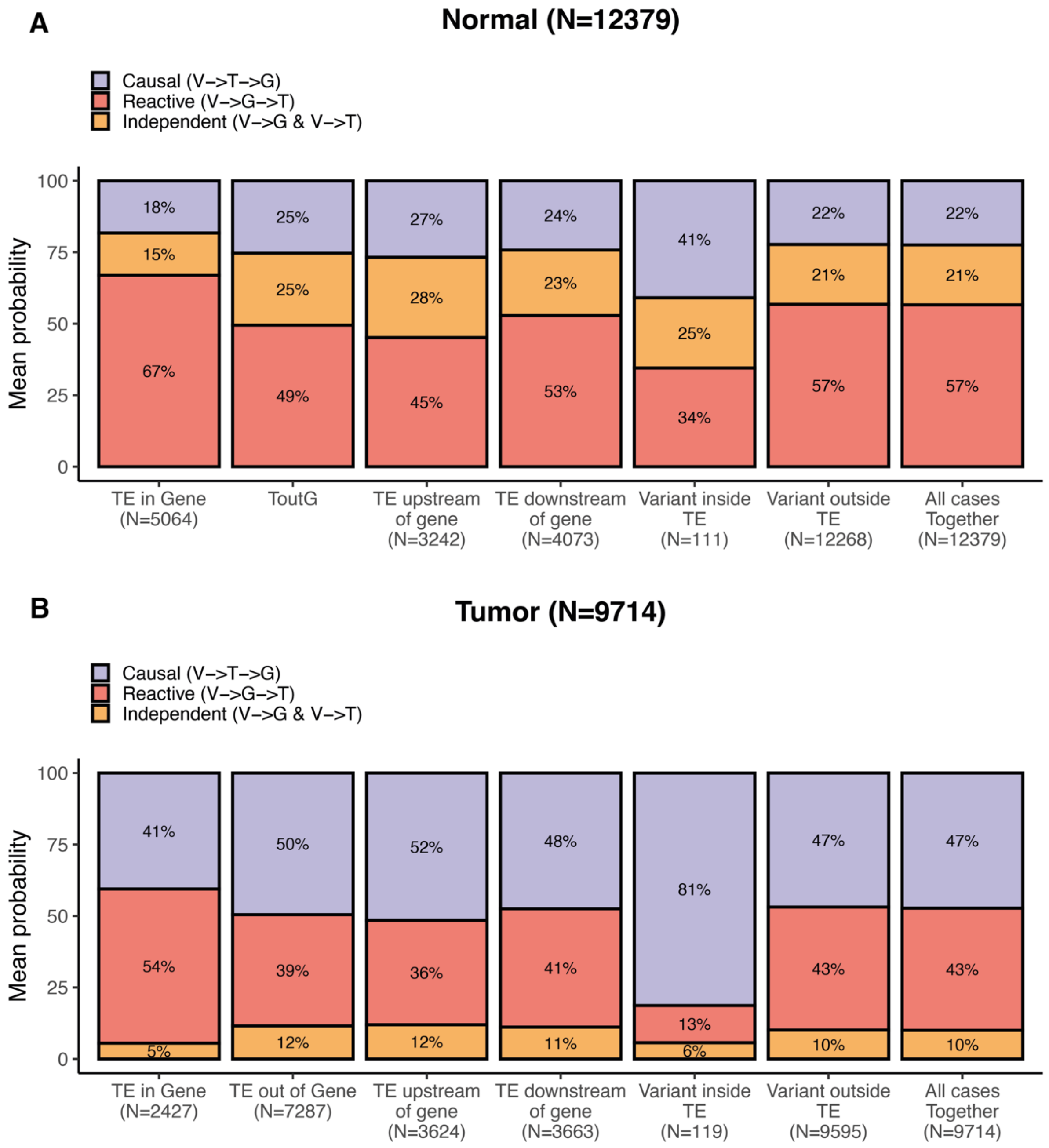


**Supplementary figure 19 | Causal relationships depending on the genomic position of the TE in respect to the gene.** In each case, we worked out the percentages by averaging the posteriors given by the Bayesian networks across all the triplets falling in each of the categories. We observe that TEs inside genes or downstream of genes tend to react to gene expression whereas TEs outside genes or when the eQTL variant is within the TE sequence, TEs are most likely causal for changes in gene expression.

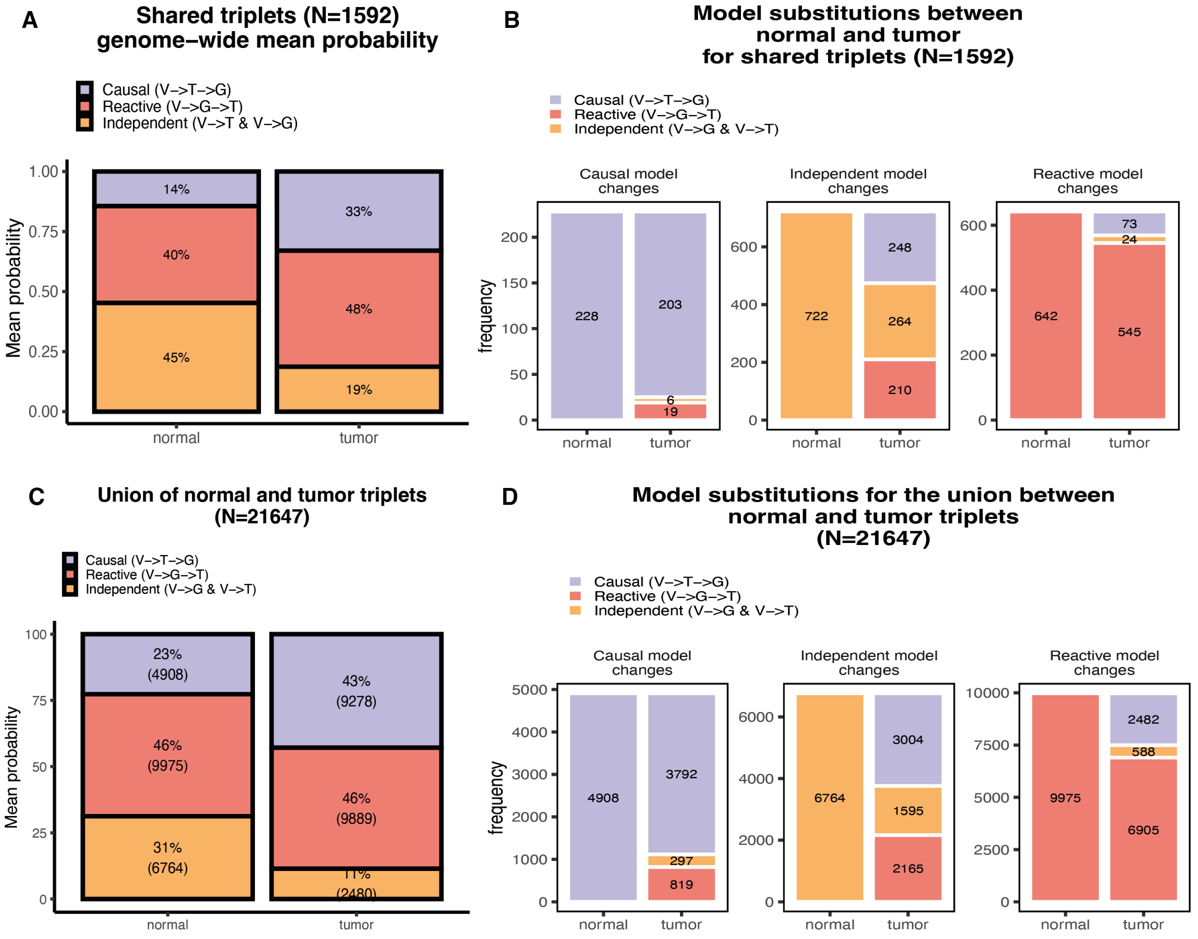


**Supplementary figure 20 | Model shifts between normal and tumor.** Model shifts for shared triplets between normal and tumor (A-B) where (A) represent the percentage of shared triplets tested that show either a higher posterior probability for the causal, reactive or independent model in tumor and normal. (**B**) Represents the model shifts from normal to tumor. (C-D) represent the same as A and B but for the union of normal and tumor triplets. In both shared and union, we observe an increase of the causal model in tumor. Independent models in normal and to a smaller extent reactive models are shifting for a causal model in tumor.

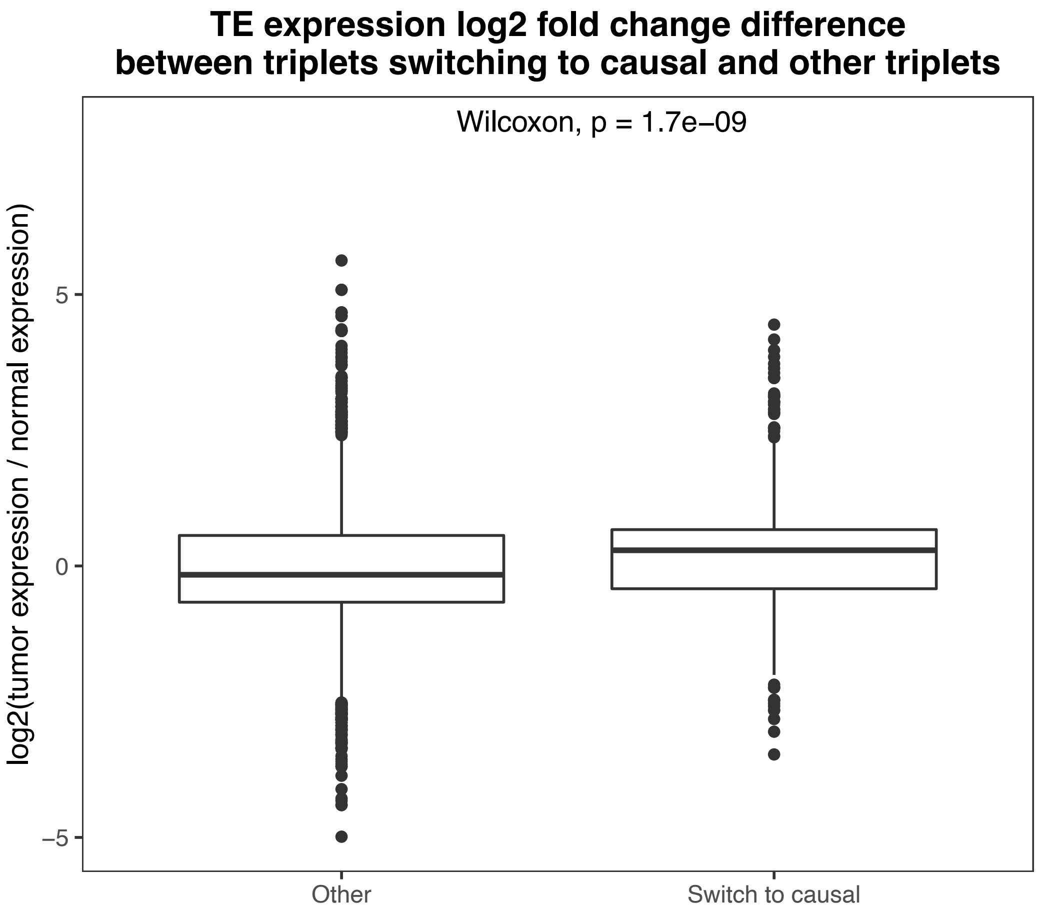


**Supplementary figure 21 | log2 fold change of TEs switching to causal versus TEs that do not switch or switch but not to causal.** We observe that TEs switching to causal in tumor are significantly more upregulated compared to TEs not switching or switching but not to causal, indicating that this upregulation could explain some of the cases where TEs switch to causal in tumor.
